# Amyloid-PET of the white matter: relationship to free water, fiber integrity, and cognition in patients with dementia and small vessel disease

**DOI:** 10.1101/2021.12.17.473211

**Authors:** Julie Ottoy, Miracle Ozzoude, Katherine Zukotynski, Min Su Kang, Sabrina Adamo, Christopher Scott, Joel Ramirez, Walter Swardfager, Benjamin Lam, Aparna Bhan, Parisa Mojiri, Alex Kiss, Stephen Strother, Christian Bocti, Michael Borrie, Howard Chertkow, Richard Frayne, Robin Hsiung, Robert Laforce, Michael D. Noseworthy, Frank S. Prato, Demetrios J. Sahlas, Eric E. Smith, Phillip H. Kuo, Jordan A. Chad, Ofer Pasternak, Vesna Sossi, Alexander Thiel, Jean-Paul Soucy, Jean-Claude Tardif, Sandra E. Black, Maged Goubran, the Medical Imaging Trials Network of Canada (MITNEC) and Alzheimer’s Disease Neuroimaging Initiative (ADNI)

**Author notes:** Corresponding author: Maged Goubran, Ph.D., 2075 Bayview Avenue, M6 East RM 706 Toronto, Canada, M4N 3M5. These authors contributed equally to this work (Julie Ottoy, Miracle Ozzoude). These authors contributed equally to this work (Sandra E. Black, Maged Goubran). Data used in preparation of this article were obtained, in part, from the Alzheimer’s Disease Neuroimaging Initiative (ADNI) database (<u>adni.loni.usc.edu</u>). As such, the investigators within the ADNI contributed to the design and implementation of ADNI and/or provided data but did not participate in analysis or writing of this report. A complete listing of ADNI investigators can be found at: http://adni.loni.usc.edu/wp-content/uploads/how_to_apply/ADNI_Acknowledgement_List.pdf.

## Abstract

White matter (WM) injury is frequently observed along with dementia. Positron emission tomography with amyloid-ligands (Aβ-PET) recently gained interest for detecting WM injury. Yet, little is understood about the origin of the altered Aβ-PET signal in WM regions. Here, we investigated the relative contributions of diffusion MRI-based microstructural alterations, including free water and tissue-specific properties, to Aβ-PET in WM and to cognition. We included a unique cohort of 115 participants covering the spectrum of low-to-severe white matter hyperintensity (WMH) burden and cognitively normal to dementia. We applied a bi-tensor diffusion-MRI model that differentiates between (i) the extracellular WM compartment (represented via free water), and (ii) the fiber-specific compartment (via free water-adjusted fractional anisotropy [FA]). We observed that, in regions of WMH, a decrease in Aβ-PET related most closely to higher free water and higher WMH volume. In contrast, in normal-appearing WM, an increase in Aβ-PET related more closely to higher cortical Aβ (together with lower free water-adjusted FA). In relation to cognitive impairment, we observed a closer relationship with higher free water than with either free water-adjusted FA or WM PET. Our findings support free water and Aβ-PET as markers of WM abnormalities in patients with mixed dementia, and contribute to a better understanding of processes giving rise to the WM PET signal.

## 1. Introduction

^18^F-Florbetapir (^18^F-AV45) is one of the most commonly used positron emission tomography (PET) ligands to detect and quantify amyloid-β (Aβ) burden in the cortex of patients with Alzheimer’s disease (AD). Specifically, Aβ-ligands are thought to target the cross beta-sheet structure of the insoluble Aβ fibrils.^1^ In addition to targeting fibrils in the cortex, these ligands are lipophilic in nature and show high retention in the white matter (WM).^1–3^ It is hypothesized that their high WM uptake is due to a similar beta-sheet structure of myelin basic protein^4^ and that microstructural damage or demyelination of WM tracts, previously observed in AD,^5^ results in reduced ligand uptake.^6^ Furthermore, while some reported the presence of Aβ in the WM,^7–9^ others did not observe such specific binding of the Aβ-ligand to WM tissue sections and attributed high WM retention to slow WM kinetics.^3, 10, 11^ Further complexity of interpreting the WM PET signal is added through the existence of localized reduced blood-to-tissue transport, hypoperfusion of lesioned WM areas, and ligand trapping in enlarged perivascular spaces, particularly in patients with concomitant small vessel disease (SVD).^12, 13^ Taken together, the molecular basis of the Aβ-ligand binding to WM remains poorly understood, hindering the understanding and interpretation of Aβ-PET scans, especially in mixed disease cohorts.

Diffusion MRI (dMRI) is a useful imaging modality to study WM microstructure *in-vivo* and may help to better understand the binding dynamics of Aβ-ligands to WM.^14, 15^ Parameters derived from diffusion tensor imaging (DTI) modeling of dMRI data include fractional anisotropy (FA) and mean diffusivity (MD), commonly used to infer changes in fiber integrity.^16^ Prior studies have shown lower FA and higher MD in areas of WM degradation such as those frequently observed as white matter hyperintensities (WMH) on Fluid Attenuated Inversion Recovery (FLAIR) MRI.^17, 18^ In addition, some found a link between lower FA and lower PET signal in the WM; thus supporting WM Aβ-PET as a marker of microstructural damage to myelin and/or axons.^14, 19^ However, this theory may be oversimplistic given that *single*-tensor DTI metrics (such as FA) are highly contaminated by signals from both adjacent cerebrospinal fluid (CSF) and freely diffusing water in the extracellular space surrounding the WM fiber tracts.^20, 21^ As such, conventional DTI metrics can hardly differentiate between degenerative and vascular/neuroinflammatory changes underlying altered Aβ-PET uptake in the WM. This bias will be exemplified in subjects with pronounced atrophy, enlarged CSF spaces, and vasogenic edema;^22, 23^ all of which are encountered in the more general AD population with often co-existent SVD pathology.^24^

Recently, a *bi-*tensor (two-compartment) model of dMRI was proposed to isolate the extracellular free water (FW) compartment from the fiber-specific (WM fibers) compartment within every image voxel.^25, 26^ In the human brain, FW is found as blood, interstitial fluid and CSF within ventricles and surrounding brain parenchyma, while it also accumulates as a form of edema in the extracellular spaces of structures distal from CSF such as deep WM.^25^ Changes in FW may indicate extracellular processes including vascular damage and inflammation. On the other hand, changes in FW-adjusted DTI metrics (e.g., FW-adjusted FA) may be closer related to microstructural tissue damage, i.e. demyelination/axonal degeneration and fiber integrity.^25^

To our knowledge, no studies have yet investigated the link between Aβ-PET uptake in the WM and WM microstructure using multi-compartment diffusion modeling, specifically FW imaging. Particularly, such research is lacking in subjects with mixed Aβ and moderate-to-severe WM lesion burden. Therefore, our study aims to investigate the relative contributions of FW and tissue-specific microstructural changes to the ^18^F-AV45 PET signal, both in regions of WMH and normal-appearing WM (NAWM), in a cohort enriched for high WMH burden. Our secondary aim is to investigate the microstructural properties and ^18^F-AV45 PET of the WM in relation to cognition. We hypothesize that lower PET signal within MRI-visible WM lesions closely relates to higher FW, which in turn, relates to cognitive impairment. Such research is imperative for more accurate interpretation of the Aβ-PET scan with potential as a future marker of WM abnormalities.

## 2. Material & Methods

### 2.1 Participants

The study included a total of 115 participants. Fifty-eight participants were recruited from dementia/stroke-prevention clinics as part of the multicenter C6 project of the Medical Imaging Network of Canada (MITNEC-C6) across seven participating sites. These participants showed moderate-to-severe SVD burden quantified as Fazekas-score>2 and confluent periventricular WMH volumes >10cc ^27^ [median(IQR): 31.1(22.4)cc]. Further inclusion criteria included: 1) clinical diagnosis of early AD or amnestic, non-amnestic single or multi-domain mild cognitive impairment (MCI/early AD, *N*=41),^28, 29^ or minor subcortical lacunar infarct/TIA (*N*=17); 2) age≥60y; 3) education>8y; and 4) MMSE≥20. Exclusion criteria included 1) major psychiatric disorder in the past 5y; 2) history of substance abuse in the past 2y; and 3) serious/chronic systemic or neurological illness (other than AD). Additionally, fifty-seven cognitively normal (CN) and patients with MCI were included from the Alzheimer’s Disease Neuroimaging Initiative (ADNI-2) database with low-to-moderate WMH volumes [median(IQR): 5.8(9.2)cc]. Inclusion criteria included: 1) CN or MCI based on the absence (CN) or presence (MCI) of memory complaints, MMSE≥24, and Clinical Dementia Rating of 0 (CN) or 0.5 (MCI); 2) age≥60y; and 3) education>8y. More details on inclusion/exclusion criteria in MITNEC-C6 and ADNI are described in **Supplementary Table 1** and at www.adni-info.org. **Table 1** describes demographics for both groups (i.e., ADNI-2 and MITNEC-C6, further referred to as the *low* and *high* WMH group, respectively). Both groups were matched for vascular risk factors including hypertension, pulse pressure, body mass index, sex, and smoking status.^30^ All participants underwent ^18^F-AV45 PET, 3T MRI including 3D T1, FLAIR, and dMRI, and a neuropsychological assessment. MRI–PET and MRI–neuropsychology acquisitions were acquired in close proximity (54±80 and 27±78 days ± standard deviation, respectively).

**Table 1.**
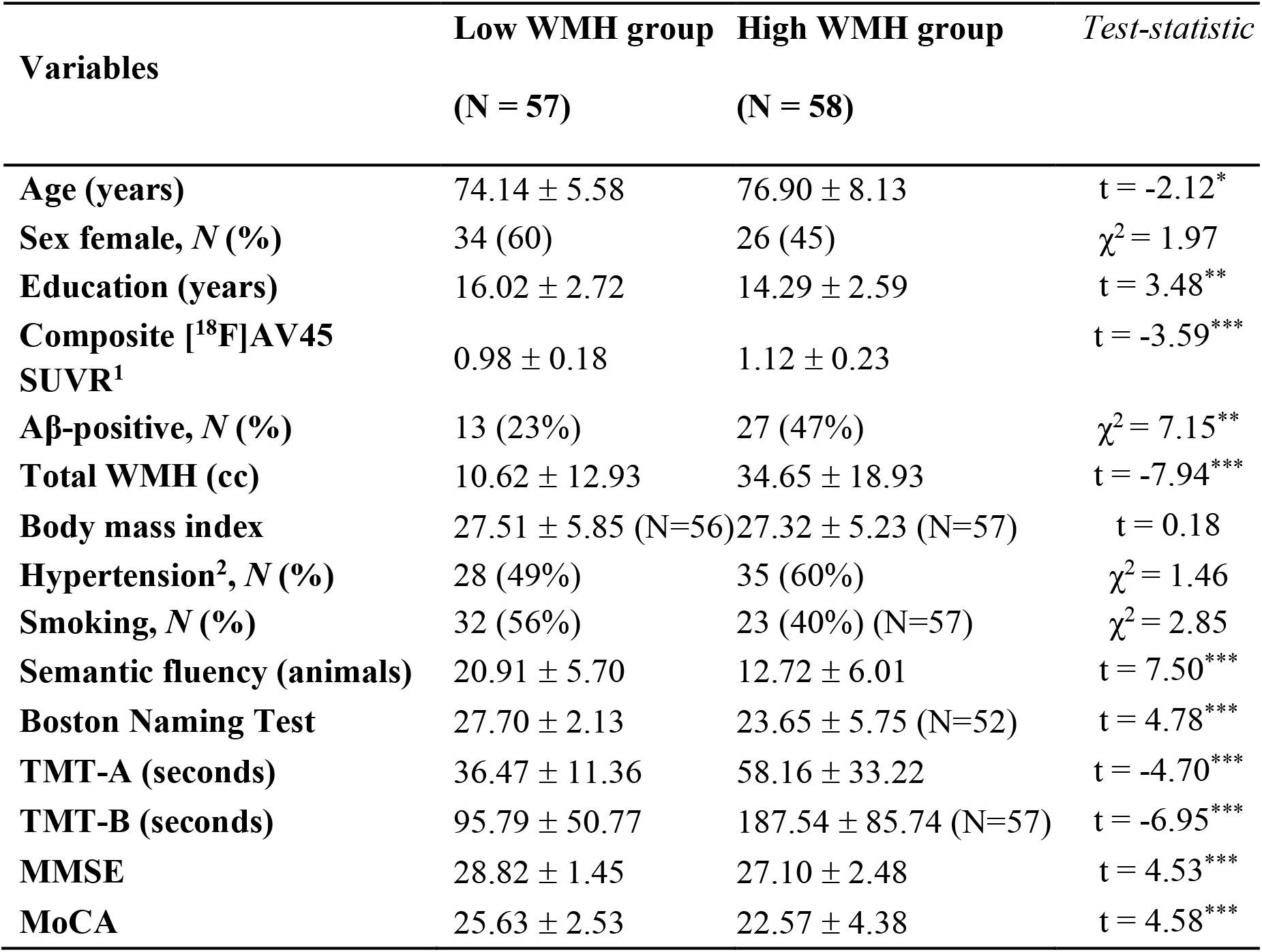
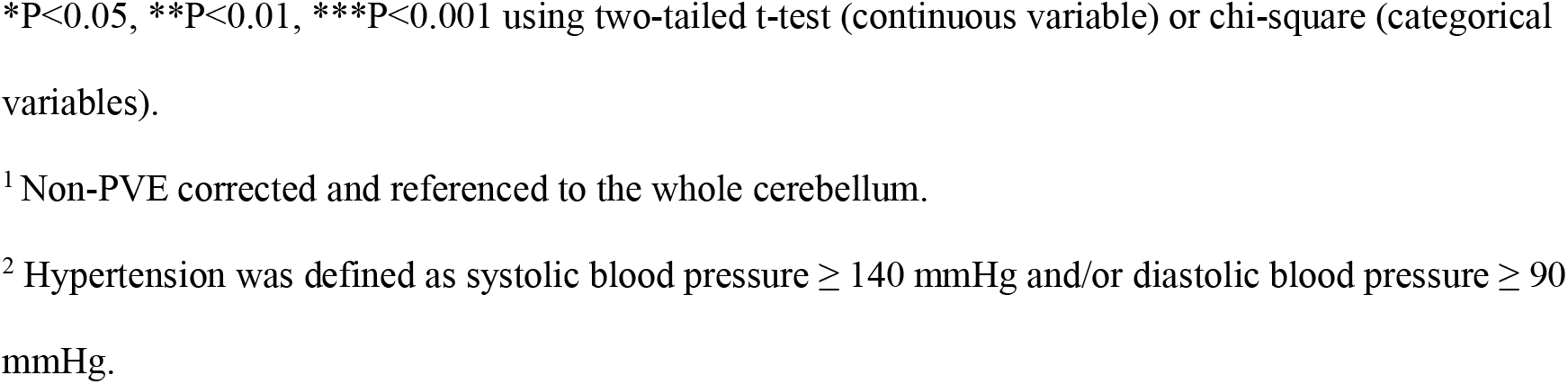
Demographics. Demographics within the low and high WMH groups. All values are indicated as mean ± standard deviation. Abbreviations: MMSE, Mini-Mental State Examination; MoCA, Montreal Cognitive Assessment; TMT, Trail Making Test; WMH, white matter hyperintensity volumes.

| **Variables** | **Low WMH group**<br>**(N = 57)** | **High WMH group**<br>**(N = 58)** | **Test-statistic** |
| --- | --- | --- | --- |
| **Age (years)** | 74.14 ± 5.58 | 76.90 ± 8.13 | t = -2.12* |
| **Sex female, N (%)** | 34 (60) | 26 (45) | $\chi^2 = 1.97$ |
| **Education (years)** | 16.02 ± 2.72 | 14.29 ± 2.59 | t = 3.48** |
| **Composite [$^{18}\text{F}$]AV45**<br>**SUVR<sup>1</sup>** | 0.98 ± 0.18 | 1.12 ± 0.23 | t = -3.59*** |
| **Aβ-positive, N (%)** | 13 (23%) | 27 (47%) | $\chi^2 = 7.15^{**}$ |
| **Total WMH (cc)** | 10.62 ± 12.93 | 34.65 ± 18.93 | t = -7.94*** |
| **Body mass index** | 27.51 ± 5.85 (N=56) | 27.32 ± 5.23 (N=57) | t = 0.18 |
| **Hypertension<sup>2</sup>, N (%)** | 28 (49%) | 35 (60%) | $\chi^2 = 1.46$ |
| **Smoking, N (%)** | 32 (56%) | 23 (40%) (N=57) | $\chi^2 = 2.85$ |
| **Semantic fluency (animals)** | 20.91 ± 5.70 | 12.72 ± 6.01 | t = 7.50*** |
| **Boston Naming Test** | 27.70 ± 2.13 | 23.65 ± 5.75 (N=52) | t = 4.78*** |
| **TMT-A (seconds)** | 36.47 ± 11.36 | 58.16 ± 33.22 | t = -4.70*** |
| **TMT-B (seconds)** | 95.79 ± 50.77 | 187.54 ± 85.74 (N=57) | t = -6.95*** |
| **MMSE** | 28.82 ± 1.45 | 27.10 ± 2.48 | t = 4.53*** |
| **MoCA** | 25.63 ± 2.53 | 22.57 ± 4.38 | t = 4.58*** |

\*P<0.05, \*\*P<0.01, \*\*\*P<0.001 using two-tailed t-test (continuous variable) or chi-square (categorical variables).
<sup>1</sup> Non-PVE corrected and referenced to the whole cerebellum.
<sup>2</sup> Hypertension was defined as systolic blood pressure $\geq 140$ mmHg and/or diastolic blood pressure $\geq 90$ mmHg.

### 2.2 Standard Protocol Approvals, Registrations, and Patient Consents

All imaging acquisition protocols were standardized across vendor platforms and compatible with ADNI-2 (see acquisition parameters in **Supplementary Table 2)**.^31^ Image quality control, archiving of clinical and imaging data, and access to pipelines were performed at the Sunnybrook Research Institute through a common database. All participants provided written informed consent. This study was conducted in full conformance with the principles of the “Declaration of Helsinki”. The institutional review board at the Sunnybrook Health Sciences Centre approved this study (Approval No. 2989).

Regarding ADNI data; ADNI was launched in 2003 as a public-private partnership. The primary goal of ADNI was to test whether MRI, PET, other biological markers, and clinical data can be combined to measure the progression of MCI/AD [see www.adni-info.org].

### 2.3 Cognitive Screening Tests

Cognitive screening tests used for this study are previously described^30^ and included processing speed and executive function (attention switching and working memory) measured with the Trail Making Test (TMT) parts A (TMT-A; *N*=115) and B (TMT-B; *N*=114), semantic fluency with animal naming (*N*=115), language with Boston Naming Test (BNT; *N*=109), and global cognition with the Montreal Cognitive Assessment (MoCA) and the Mini-Mental State Examination (MMSE) (*N*=115). TMT scores were log-transformed to achieve a normal distribution.

### 2.4 Structural MRI

MRI was performed using acquisition protocols that were standardized across vendor platforms and compatible with the ADNI-2 protocol (see acquisition parameters in **Supplementary Table 2)**.^31^ The T1-weighted (T1w) MRI images were processed with FreeSurfer v6.0 using an in-house modified pipeline for subjects with SVD.^32^ Subcortical lacunar infarcts were masked out before WMH segmentation.^33^ WMH were delineated based on our automated segmentation tool HyperMapper.^34^ More details on MRI processing are described in **Supplementary methods S1**. Total WMH volume was normalized by the total intracranial volume (ICV), and log-transformed. Total ICV was estimated using a skull-stripping deep learning network developed in-house that is robust to vascular lesions and atrophy.^35^

The NAWM was segmented in native space by subtracting the binarized WMH regions from the binarized whole WM defined by the Desikan-Killiany-Tourville atlas. To limit partial volume effects (PVE), we 1) did not include ADNI subjects with WMH <1cc (no MITNEC-C6 subjects fulfilled this criterion), 2) used a stricter HyperMapper threshold on segmentation probabilities (0.4-0.5) to avoid overestimation of WMH,^34^ and 3) performed erosion on the NAWM masks by 2×2×2mm^3^ to eliminate spill-effects from neighboring tissue including cortex and CSF. **Figure 1A** highlights WMH and NAWM delineations in an example subject. We also tested for different NAWM erosion kernels including 1×1×1mm^3^ and 3×3×3mm^3^.

**Figure 1.**
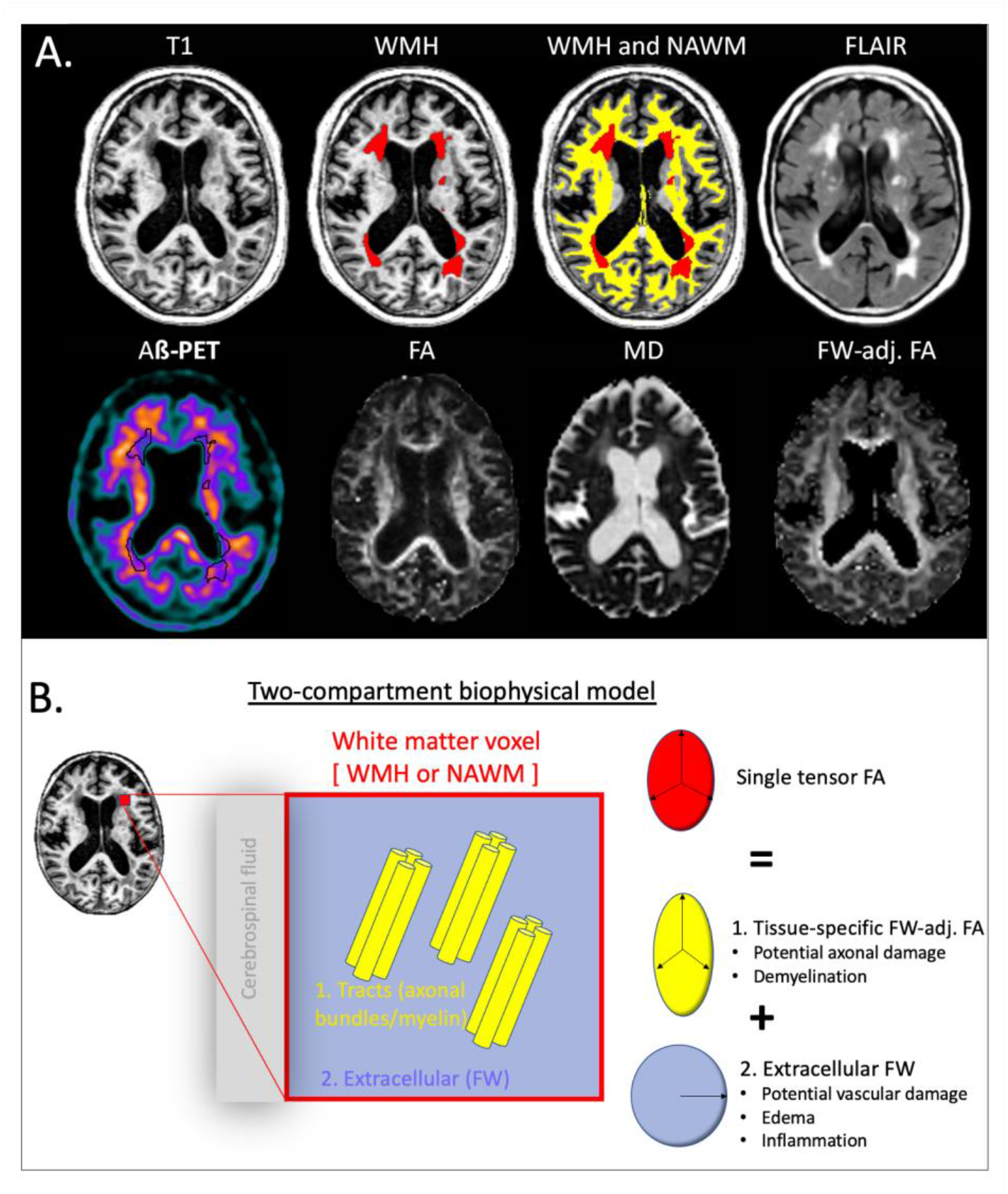
Overview of methods. A) Imaging acquisitions and segmentations in a representative subject. (Upper panel, left to right:) T1-w MRI image, T1w image with WMH delineation (red), T1w image with delineation of both WMH (red) and NAWM (yellow), FLAIR image. (Lower panel, left to right:) ^18^F-AV45 PET, FA, MD, and FW-adjusted FA maps showing altered signal in the regions of WMH. B) Schematic representation of the two-compartment biophysical DTI model. Each brain voxel (red color) of the NAWM or WMH can be separated into two compartments: an extracellular (FW; purple color) and a fiber-specific (FW-adjusted FA; yellow color) compartment. The ‘adjusted’ tissue diffusion ellipsoid is more prolate (yellow tensor) after being separated from the isotropic diffusion ellipsoid (purple tensor). Abbreviations: FA, fractional anisotropy; FW, free water; NAWM, normal-appearing white matter; WMH, white matter hyperintensities.

### 2.5 Diffusion tensor imaging

Raw ADNI data was downloaded from the ADNI-2 database. Acquisition parameters included 256×256 matrix size, 1.4×1.4×2.7mm^3^ voxel size, 9000ms TR, and 9min scan time. A total of 41 diffusion encoding directions (*b*=1000s/mm^2^) and five *b*=0s/mm^2^ were acquired. More details can be found at http://adni.loni.usc.edu/wp-content/uploads/2010/05/ADNI2_GE_3T_22.0_T2.pdf. In MITNEC-C6, dMRI scans were acquired using a 128×128 matrix size, 2mm isotropic voxels, TR ranging between 9400 and 10000ms and 30 or 32 directions depending on the vendor with *b*=1000s/mm^2^ and two *b*=0s/mm^2^. These acquisition protocols were standardized across vendor platforms (see **Supplementary Table 2)**.^31^ Raw data from both cohorts were processed with the same imaging pipeline. The preprocessing steps to obtain corrected dMRI data and DTI scalar maps (FA and MD) were in line with Nir et al.^36^ and described in more detail in **Supplementary methods S2**.

For FW mapping, the eddy current and motion-corrected dMRI data were fitted to a two-compartment diffusion model in each voxel, separating the FW compartment from the non-FW tissue compartment (**Figure 1B**).^25^ Specifically, an FW map represents the fractional volume (ranging from 0 to 1) of freely diffusing extracellular water with a fixed isotropic diffusivity of 3×10^−3^ mm^2^/s (diffusion coefficient of water at body temperature). The FW-corrected FA represents the FA signal following removal of the FW signal and is expected to be more specific to axonal and myelination changes than FA alone (**Figure 1B**).^20^ Subsequently, the mean FA, MD, FW, and FW-adjusted FA were extracted in the whole brain, WMH, and NAWM. **Figure 1A** shows an example of a T1w image with corresponding DTI-based scalar maps.

FW correlated strongly (Pearson’s r > 0.5) with both FA and MD in WMH and with FA in NAWM, but not with FW-adjusted FA indicating that the FW effect was indeed removed from FW-adjusted FA (**Supplementary Figure 1**).

### 2.6 Amyloid-PET imaging

Details on acquisition and processing of ^18^F-AV45 scans are described in **Supplementary methods S3**. The raw data of both cohorts were processed with the same imaging pipeline.^30^ Standardized uptake value ratios (SUVR) were generated in the WMH and NAWM masks within the native T1w space and referenced to the whole cerebellum after removing T1-hypointense regions corresponding to cerebellar WM abnormalities with FreeSurfer. Previous work indicated the relevance of including WM in the reference region for AV45 quantification in dementia cohorts.^2^,^37, 38^ A ‘global’ Aβ SUVR value was derived based on the AD-signature regions (volume-weighted average of the frontal, parietal, temporal, and cingulate regions)^39^ and corrected for PVE.

### 2.7 Statistical analyses

Statistics were performed using Python v3.7, R v4.0, and PROCESS v3.5 in SPSS. All continuous metrics were z-scored. Model-based measures were reported as effect estimates and 95% confidence intervals (CI) based on bootstrapping with 1,000 replications. A paired t-test was used to detect significant differences in SUVR or DTI metrics between NAWM and WMH. The analyses described below were investigated across all subjects, as well as within the high and low WMH groups separately.

#### 2.7.1 Relationship between ^18^F-AV45 SUVR and DTI metrics

Linear regressions were used to assess the associations between each of the DTI metrics (independent variable) and SUVR (dependent variable) in NAWM or WMH, adjusted for continuous age and education, and sex. The regressions evaluated in the WMH were additionally adjusted for global cortical Aβ SUVR and for WMH volume. The regressions evaluated in the NAWM were adjusted for global cortical Aβ SUVR; WMH volume was not significantly associated with NAWM SUVR and thus omitted as covariate (P>0.05).

Partial least square (PLS) regression analyses were performed to further investigate how the DTI, demographical (age, sex, education) and imaging (WMH volume, cortical Aβ SUVR) variables covary together in predicting SUVR in the WMH or NAWM. PLS analysis assures that, when a set of predictors is large and closely related to one another (FA, FW-adjusted FA, MD, and FW), the optimum subset of predictors can be selected by extracting orthogonal components that explain as much as possible covariance between X and Y. The number of components was determined based on the root-mean-square error derived from a ten-fold cross-validation with five repeats (**Supplementary Figure 2**).

#### 2.7.2 Relationship with cognitive impairment

Linear regression analysis was employed to assess the associations between DTI metrics in WMH or NAWM (FW and FW-adjusted FA; independent variable) and cognitive scores (dependent variable), adjusted for age, sex, education, WMH volume, and global cortical Aβ SUVR. In addition, the relationship between SUVR in the WMH or NAWM and cognition was investigated with mediation analysis using FW or FW-adjusted FA as the mediators, adjusted for the aforementioned covariates. Bias-corrected bootstrapping with 5,000 replications and a 95% CI was performed for estimation of (in)direct and total effects.

#### 2.7.3 Linear mixed-effect models

To investigate the effects of age or sex on ^18^F-AV45 SUVR in the WM, we employed linear mixed effect models with an interaction term between age or sex and WM region (NAWM and WMH) and subject as a random factor.

### 2.8 Data and materials availability

All data associated with this study are available in the main text or supplementary materials. All the imaging data can be shared upon request with a proposal and under a data transfer agreement.

## 3. Results

### 3.1 Demographics

The high WMH group (*N*=58) was significantly older and less educated compared to the low WMH group (*N*=57). Additionally, they had significantly higher cortical Aβ load and lower cognitive scores (**Table 1**). The proportion of Aβ-positive subjects was 22.8% and 46.6% in the low and high WMH group, respectively.

### 3.2 ^18^F-AV45 SUVR and DTI metrics are altered in WMH compared to NAWM

Regions of WMH showed significantly lower SUVR compared to NAWM (∼14% reduction, t=25.08, P<0.0001) (**Figure 2**). In addition, regions of WMH showed significantly higher FW (t=-27.11), higher MD (t=-24.37), lower FA (t=32.77), and lower FW-adjusted FA (t=12.26) compared to NAWM (all P<0.0001; **Figure 2**).

**Figure 2.**
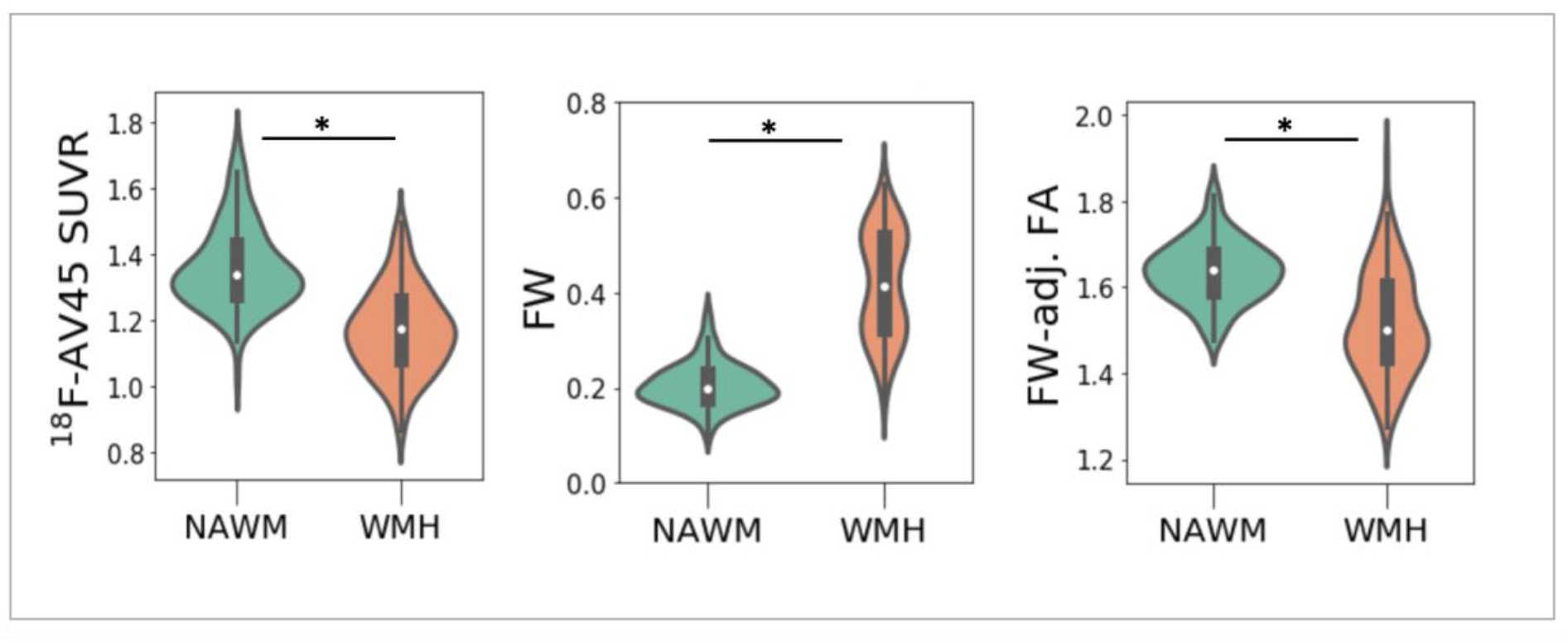
Differences in ^18^F-AV45 SUVR and DTI metrics between regions of WMH and NAWM. Violin plots representing the group differences in SUVR (left panel), FW (middle panel), and FW-adjusted FA (normalized to whole brain value; right panel) in the WMH (orange) vs. NAWM (green) across all subjects. Paired t-test showed significant differences between WMH and NAWM at *P<0.0001 for all metrics. Specifically, SUVR and FW-adjusted FA were significantly lower and FW higher in WMH compared to NAWM. Abbreviations: FA, fractional anisotropy; FW, free water; NAWM, normal appearing white matter; SUVR, standardized uptake value ratio; WMH, white matter hyperintensities.

### 3.3 ^18^F-AV45 SUVR relates to FW in WMH and to FW-adjusted FA in NAWM

#### 3.3.1 Regression analyses of diffusion metric predicting WM SUVR

In regions of WMH, lower SUVR was strongly associated with higher FW both across all subjects (β=-0.36, P=0.005; 95%CI_bootstrap_[-0.63,-0.10]) (**Figure 3A**-left) and in the high WMH group separately (β=-0.33, P=0.019; 95%CI_bootstrap_[-0.55,-0.06]), independent of cortical Aβ. In addition, in regions of WMH, lower SUVR was associated with lower FA (β=+0.24, P=0.046; 95%CI_bootstrap_[+0.008,+0.47]) and showed a trend with higher MD (P=0.064; 95%CI_bootstrap_[-0.44,+0.003]) across all subjects, but not with the fiber-specific metric FW-adjusted FA. Taken together, these results indicate that, amongst the DTI metrics, FW was most closely associated with ^18^F-AV45 SUVR in regions of WMH.

**Figure 3.**
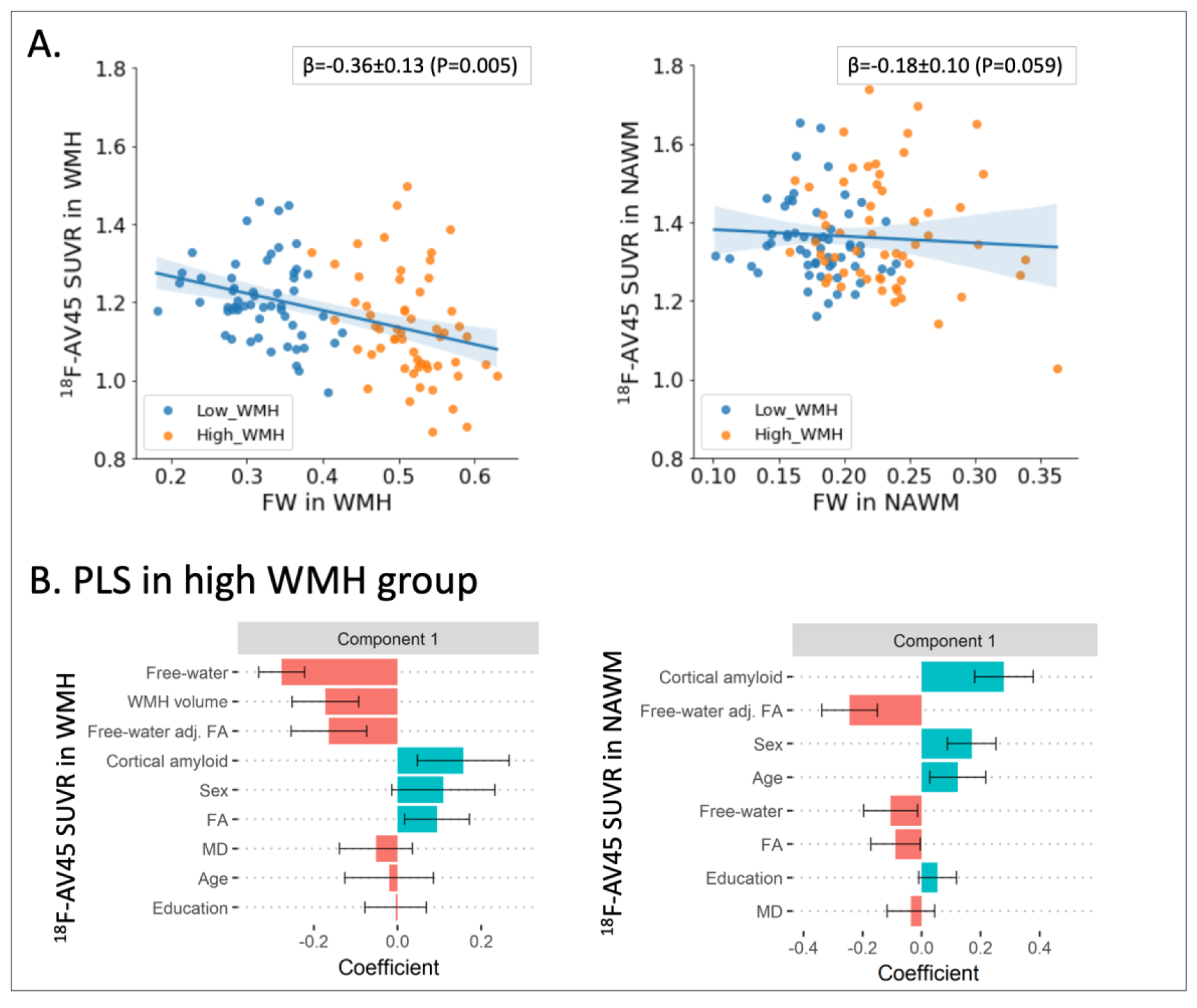
Relationship between FW and ^18^F-AV45 SUVR. A) Increased FW is associated with reduced SUVR in WMH (left) but not in NAWM (right). Data points are colored based on whether the subject belongs to the low (blue) or high (orange) WMH group. B) PLS analysis showing how DTI metrics covary together in predicting SUVR in regions of the WM. The plots represent the contribution of the loadings to the first component of PLS analysis explaining most of the variance in SUVR in WMH (left panel; component 1 ∼ 24%, component 2 [data not shown] ∼ 2%) and NAWM (right panel; component 1 ∼ 31%, component 2 [data not shown] ∼ 2%) in subjects belonging to the high WMH group. Predictors include DTI metrics (FA, MD, FW, and FW-adjusted FA), demographics (age, female sex, education), and imaging variables (WMH volume and cortical Aβ SUVR). Predictors are ordered based on the absolute value of the loading: FW and WMH volume had the most influence on signal in the WMH while cortical Aβ and FW-adjusted FA had the most influence on signal in the NAWM. Error bars represent 95%CI based on bootstrapping with 5,000 repetitions. PLS results across all subjects and within the low WMH group are represented in **Supplementary Figure 4**. Abbreviations: FW, free water; MD, mean diffusivity; NAWM, normal appearing white matter; SUVR, standardized uptake value ratio; WMH, white matter hyperintensities.

In the NAWM, SUVR was inversely associated with FW-adjusted FA in the high WMH group (β=-0.32, P=0.008; 95%CI_bootstrap_[-0.53,-0.07]) and showed a trend across all subjects (P=0.077; 95%CI_bootstrap_[-0.32,+0.05]), independent of cortical Aβ. Different erosion kernels for the NAWM mask obtained similar results (**Supplementary Figure 3**). In addition, there was a trend towards an association between lower SUVR and higher FW across all subjects (P=0.059; 95%CI_bootstrap_[-0.38,+0.05]; **Figure 3A**right), while no associations were found with MD or FA in the NAWM. Taken together, these results indicate that, amongst the DTI metrics, FW-adjusted FA was most closely associated with ^18^F-AV45 SUVR in the NAWM (particularly in the high WMH subgroup).

Due to the recruitment from different cohorts across all subjects and from different clinics within the high WMH group, we repeated our analyses adjusting for this factor. **Supplementary Table 3** indicates that results remained the same and that cohort/clinic was a non-significant covariate in the analyses.

#### 3.3.2 PLS analyses of diffusion metrics predicting WM SUVR

Our findings were further confirmed by PLS analysis of WM SUVR with all the DTI metrics together in the model. In regions of WMH, FW and WMH volume loaded most strongly inversely onto the first component (24% variance), while FW-adjusted FA and FA also had a significant but smaller contribution (**Figure 3B**-left; high WMH group). This may indicate that lower SUVR in regions of WMH is mainly associated with higher FW together with higher WMH volume. On the other hand, in NAWM, cortical Aβ SUVR loaded most strongly positively onto the first component (31% variance), while FW and FW-adjusted FA also had a significant but smaller contribution (**Figure 3B**-right; high WMH group). This may indicate that higher SUVR in NAWM is mainly associated with higher cortical Aβ rather than diffusion alterations. Similar results with PLS analysis were found across all subjects and within the low WMH group separately, i.e., FW, WMH volume, and FA consistently contributed to WMH SUVR, while cortical Aβ SUVR consistently contributed to NAWM SUVR (**Supplementary Figure 4**). Interestingly, while FW-adjusted FA loaded inversely onto NAWM SUVR for the high WMH group (**Figure 3B**-right), it loaded positively for the low WMH group (**Supplementary Figure 4B**).

### 3.4 Increased FW in WM is associated with cognitive impairment

In regions of WMH, higher FW was associated with worse performance on the MoCA, MMSE, language, and semantic fluency tests both across all subjects and in the high WMH group separately (**Figure 4A**; **Table 2; Supplementary Table 4** with adjustment for clinic). In fact, higher FW mediated the relationship between WMH SUVR and executive function (β_indirect_=-0.06, 95%CI_bootstrap_[-0.14,-0.01]). Conversely, lower FW-adjusted FA was significantly associated only with lower MoCA scores across all subjects.

**Figure 4.**
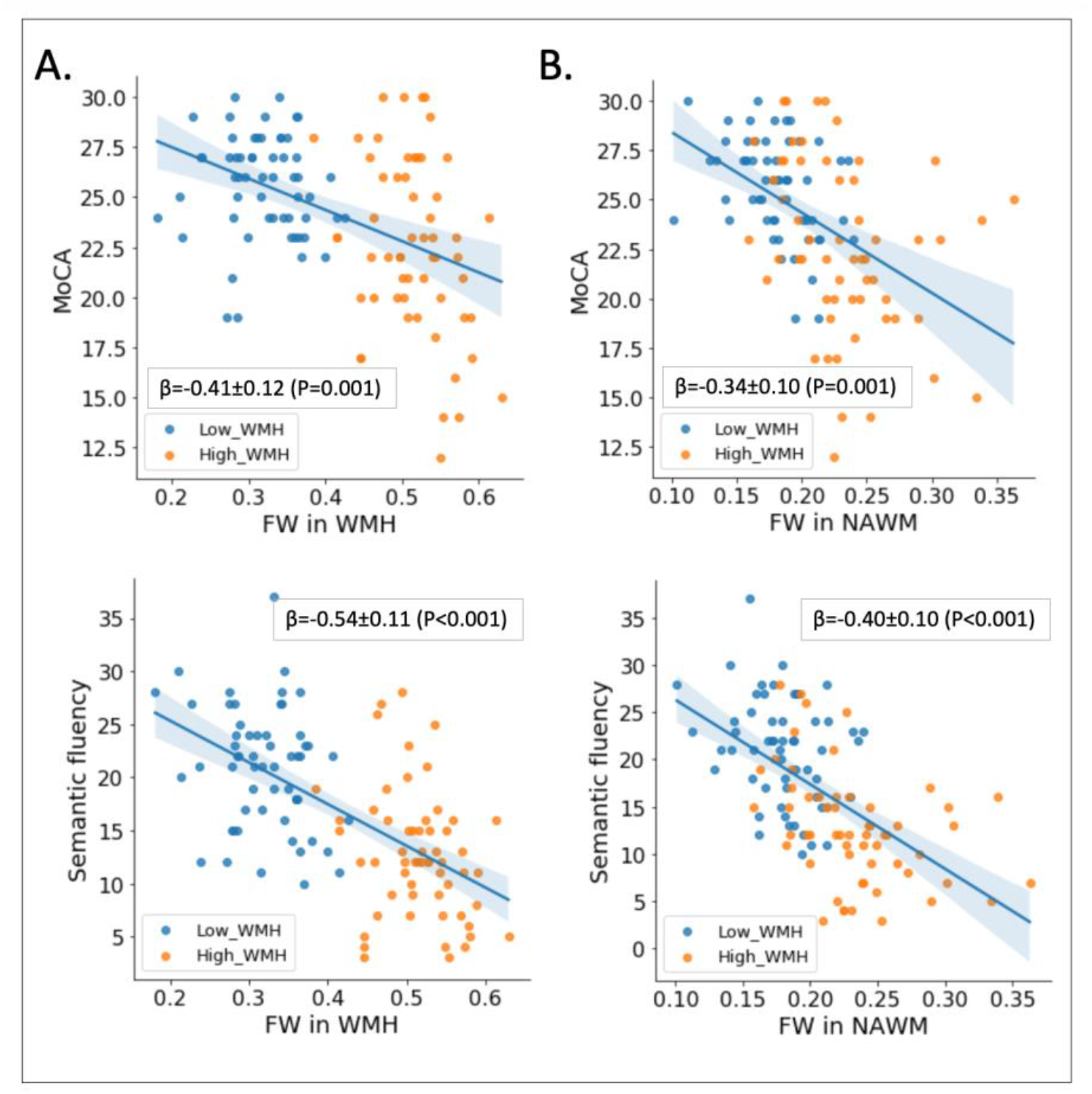
Associations between FW and cognition. Increased FW in the WMH (panel A) or NAWM (panel B) associate with cognitive impairment, including MoCA (upper row) and semantic fluency (lower row), across all subjects. Data points are colored based on whether the subject belongs to the low (blue) or high (orange) WMH group. Abbreviations: FW, free water; MoCA, Montreal Cognitive Assessment; NAWM, normal appearing white matter; WMH, white matter hyperintensities.

**Table 2.** Association between DTI metrics and cognition in regions of WMH and NAWM. The regression analyses were adjusted for age, sex, education, WMH volume, and global cortical Aβ SUVR. Results are shown separately for all subjects combined (n=115) and the high WMH subgroup (n=58). Abbreviations: bs, bootstrap (1,000 repetitions); FA, fractional anisotropy; FW, free water; MMSE, Mini-Mental State Examination; MoCA, Montreal cognitive assessment; WMH, white matter hyperintensities

| **Regions of white matter hyperintensities** |  |  |  |  |  |  |
| --- | --- | --- | --- | --- | --- | --- |
| *All subjects (n=115)* |  |  |  |  |  |  |
|  | **FW** |  |  | **FW-adjusted FA** |  |  |
| | $\beta$ | *P* | 95%CI<sub>bs</sub> | $\beta$ | *P* | 95%CI<sub>bs</sub> |
| *Semantic* | -0.54 | **<0.0001** | -0.78,-0.29 | +0.14 | **0.084** | -0.03,+0.31 |
| *Language* | -0.38 | **0.002** | -0.65,-0.10 | +0.17 | **0.050** | +0.03,+0.34 |
| *MoCA* | -0.41 | **0.001** | -0.69,-0.12 | +0.17 | **0.048** | -0.003,+0.33 |
| *MMSE* | -0.40 | **0.003** | -0.67,-0.10 | +0.14 | *0.11* | -0.02,+0.32 |
| *Speed* | +0.31 | **0.016** | +0.01,0.58 | -0.07 | *0.44* | -0.22,+0.08 |
| *Executive* | +0.35 | **0.002** | +0.10,+0.59 | -0.10 | *0.19* | -0.26,+0.05 |
| *High WMH group (n=58)* |  |  |  |  |  |  |
|  | **FW** |  |  | **FW-adjusted FA** |  |  |
| | $\beta$ | *P* | 95%CI<sub>bs</sub> | $\beta$ | *P* | 95%CI<sub>bs</sub> |
| *Semantic* | -0.26 | **0.059** | -0.54,+0.02 | +0.08 | *0.58* | -0.16,+0.32 |
| *Language* | -0.40 | **0.003** | -0.68,-0.15 | +0.23 | **0.091** | -0.05,+0.47 |
| *MoCA* | -0.34 | **0.013** | -0.61,-0.04 | +0.11 | *0.41* | -0.14,+0.39 |
| *MMSE* | -0.31 | **0.036** | -0.52,-0.03 | +0.08 | *0.61* | -0.18,+0.41 |
| *Speed* | +0.08 | *0.56* | -0.21,+0.41 | -0.11 | *0.45* | -0.33,+0.14 |
| *Executive* | +0.20 | *0.15* | -0.01,+0.48 | -0.17 | *0.20* | -0.52,+0.07 |

Region of normal-appearing white matter
| *All subjects (n=115)* |  |  |  |  |  |  |
| --- | --- | --- | --- | --- | --- | --- |
|  | FW |  |  | FW-adjusted FA |  |  |
| | $\beta$ | *P* | 95%CI<sub>bs</sub> | $\beta$ | *P* | 95%CI<sub>bs</sub> |
| *Semantic* | -0.40 | **<0.0001** | -0.61,-0.21 | +0.03 | *0.71* | -0.14,+0.22 |
| *Language* | -0.23 | **0.032** | -0.42,-0.05 | +0.18 | **0.034** | -0.03,+0.40 |
| *MoCA* | -0.34 | **0.001** | -0.60,-0.11 | +0.10 | *0.23* | -0.09,+0.31 |
| *MMSE* | -0.30 | **0.010** | -0.54,-0.09 | +0.21 | **0.020** | +0.01,+0.41 |
| *Speed* | +0.32 | **0.003** | +0.08,+0.53 | +0.05 | *0.59* | -0.12,+0.22 |
| *Executive* | +0.23 | **0.021** | +0.04,+0.42 | -0.05 | *0.49* | -0.21,+0.09 |
| *High WMH group (n=58)* |  |  |  |  |  |  |
|  | FW |  |  | FW-adjusted FA |  |  |
| | $\beta$ | *P* | 95%CI<sub>bs</sub> | $\beta$ | *P* | 95%CI<sub>bs</sub> |
| *Semantic* | -0.40 | **0.006** | -0.67,-0.09 | -0.14 | *0.35* | -0.37,+0.16 |
| *Language* | -0.27 | **0.064** | -0.56,+0.03 | +0.09 | *0.53* | -0.16,+0.43 |
| *MoCA* | -0.32 | **0.028** | -0.65,-0.01 | -0.01 | *0.93* | -0.29,+0.29 |
| *MMSE* | -0.36 | **0.018** | -0.68,-0.11 | +0.08 | 0.60 | -0.25,+0.39 |
| *Speed* | +0.24 | 0.10 | -0.07,+0.52 | +0.14 | 0.33 | -0.10,+0.45 |
| *Executive* | +0.15 | 0.30 | -0.11,+0.42 | -0.08 | 0.59 | -0.32,+0.14 |

Following cognitive tests were applied. Semantic fluency: animal naming; Language: BNT; Speed: TMT-A;
Executive function: TMT-B

Similarly, in the NAWM, higher FW was associated with worse performance on the MoCA, MMSE, language, and semantic fluency tests both across all subjects and in the high WMH group separately (**Figure 4B**; **Table 2; Supplementary Table 4** with adjustment for clinic). Conversely, lower FW-adjusted FA in the NAWM was significantly associated only with lower MMSE across all subjects.

### 3.5 Effects of age and sex on ^18^F-AV45 SUVR in the WM

We observed a significant interaction effect (P<0.0001) between age and WM regions (i.e., NAWM vs WMH) on SUVR, such that older age was associated with lower SUVR in the WMH but with higher SUVR in the NAWM (**Supplementary Figure 5**). Females showed consistently higher SUVR in both NAWM (P=0.008) and WMH (P=0.006) compared to males, with no observed interaction (**Supplementary Figure 6**).

## 4. Discussion

This study investigated the neurobiological underpinnings of Aβ-PET (^18^F-AV45) uptake in the WM using two-compartment modelling of diffusion MRI in a mixed cohort of patients with mild-to-severe SVD (manifested as WMH) and cortical Aβ. In line with current literature, we observed that SUVR in regions of WMH was significantly decreased compared to the NAWM SUVR.^14, 15, 19^ Our manuscript led to three novel observations (summary **Figure 5**): 1) In WMH, the decrease in SUVR was closely related to higher FW; 2) In NAWM, the increase in SUVR was closely related to higher cortical Aβ (together with lower FW-adjusted FA); and 3) In both WMH and NAWM, higher FW was related to cognitive impairment and WMH FW mediated the association between WMH SUVR and executive function. These findings can be interpreted as following: 1) The lower PET signal in WMH may largely reflect vascular damage, edema, inflammation, and/or fiber necrosis that led to concomitant enlargement of the extracellular space; 2) The higher PET signal in NAWM tracks with cortical Aβ and may partly reflect microstructural damage to the remaining fibers; and 3) FW may be a sensitive SVD-related biomarker in AD and mixed dementia to detect early and more subtle changes in WM microstructure.

**Figure 5.**
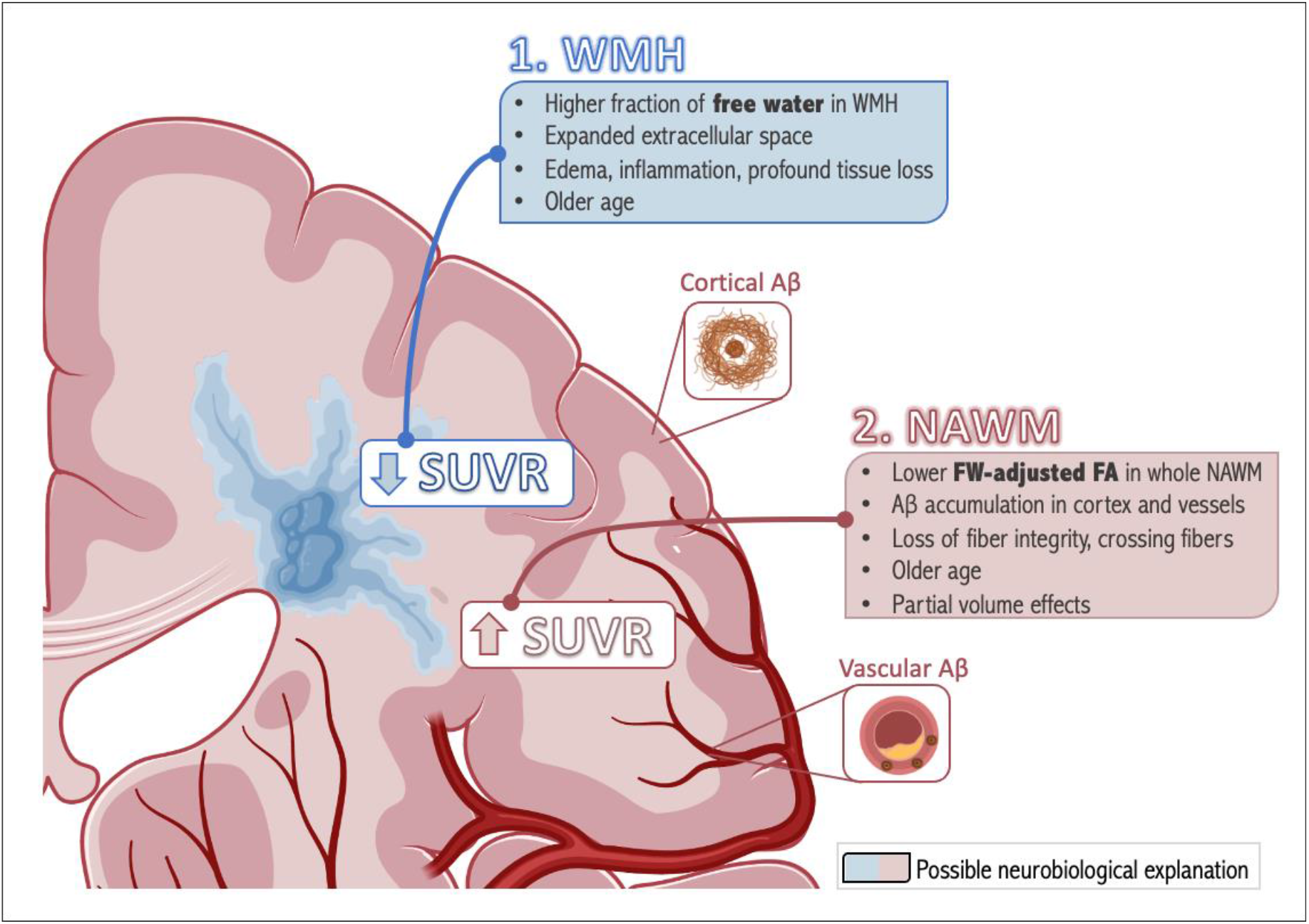
Summary of relationships between ^18^F-AV45 SUVR and DTI metrics in WM, along with potential neurobiological explanations. Abbreviations: FW-adjusted FA, free water-adjusted fractional anisotropy; NAWM, normal-appearing white matter; SUVR, standardized uptake value ratio derived from amyloid-PET; WMH, white matter hyperintensities [created with BioRender.com]

Supported by previous evidence that myelin alterations represent an early feature of aging and AD,^5^ Aβ-PET in the WM was recently suggested as a marker of local myelin integrity;^14, 15, 19^ thereby adding value to quantifying Aβ in the cortex. However, prior studies did not account for contributions of FW to the conventional DTI metrics (such as FA), ultimately limiting their interpretation and potentially overestimating the level of demyelination or axonal damage.^21, 40^ While we confirm prior studies by reporting an association between lower Aβ SUVR and lower FA (before FW adjustment) in WMH regions,^14^ we notably showed that the lower SUVR was more strongly associated with FW than with either FA or FW-adjusted FA. This lower SUVR in WMH was also associated with higher WMH volume. These findings may indicate that the PET signal within MRI-visible WM lesions is more profoundly associated with enlargement of the extracellular space (potentially of vascular or inflammatory origin, which may in part be secondary to profound tissue (myelin) loss/necrosis) than with tissue-specific compartment alterations (e.g., reduced fiber integrity or localized demyelination).^25, 41^ Several additional considerations may support this observation. First, while demyelinating fibers are an early feature in AD,^5^ their direct effects on diffusion may be less pronounced in a sample enriched for SVD.^18, 40^ Indeed, our group has previously shown that WM disease may primarily reflect chronic vasogenic edema (blood-brain barrier [BBB] leakage) or perivascular stasis (compromised circulation of interstitial fluid) induced by venous collagenosis and chronic hypoperfusion;^22, 23, 42^ which may be a trigger of downstream neuroinflammation, damage to myelin membranes with potential formation of intramyelinic fluid-filled vacuoles, and axonal (Wallerian) degeneration due to cortical neuronal injury in cases with concomitant AD pathology.^40, 41, 43^ Second, previous autopsy studies reported no PiB staining/binding to WM tracts,^3, 10^ suggesting that demyelination alone does not account for the reduced PET signal. Taken together, we speculate that reduced ^18^F-AV45 retention in MRI-visible WM lesions is at least partly reflective of increased water content.

Importantly, while the association of lower WMH SUVR with higher WMH volume and lower cortical SUVR in PLS component 1 may be suggestive of PVE, we believe that PVE cannot explain the lower WMH SUVR compared to NAWM. Indeed, we not only performed various methodological efforts to reduce PVE but also benefitted from a unique cohort with high WMH volumes less prone to PVE. Furthermore, previous studies likewise found a relationship of higher WMH volume with lower FA, higher MD, and higher interstitial fluid,^14, 17, 18, 40^ as well as lower WMH SUVR in A-compared to A+,^44^ advocating for a biological rather than PVE-related explanation underlying the lower WMH SUVR.

In NAWM, the relation of SUVR with FW-adjusted FA was stronger than with either FA or FW (**Figure 5**). This may be related to a lesser expansion of the extracellular space in NAWM. Interestingly, while lower FW-adjusted FA is typically interpreted as reduced fiber integrity, we observed a relationship between lower SUVR and higher (rather than lower) FW-adjusted FA in the whole NAWM of our high WMH subgroup. There are several potential explanations for this finding. First, higher FW-adjusted FA may represent compression of fibers by surrounding edema^45^ or loss of crossing fibers, consistent with degeneration of selected tracts as previously observed in patients with MCI or AD.^46, 47^ Second, we observed, similar to prior work, that cortical and NAWM SUVRs are positively correlated even after partial volume correction/erosion.^14, 44, 48, 49^ This may indicate that cortical Aβ SUVR is one of the drivers behind (i) increased NAWM SUVR, and (ii) decreased FW-adjusted FA (indeed, higher cortical Aβ SUVR was associated with lower FW-adjusted FA (data not shown); which may be either through direct WM injury or secondary Wallerian degeneration).^5, 50^ One potential explanation for the positive cortex-NAWM SUVR relationship, considering that NAWM was eroded, may be impaired CSF-mediated clearance and binding to diffuse Aβ, APP, or Aβ deposits in vessel walls of the WM,^7, 9, 51^ observed with aging and AD. For example, periventricular venous insufficiency resulting from stenosed or occluded vessels may interfere with interstitial cerebral fluid circulation, impairing the drainage of Aβ along the perivascular spaces and promoting Aβ deposition as plaques and around the vessels.^52^ Another potential explanation for the cortex-NAWM SUVR relationship is remaining PET-associated PVE (the strength of this relationship decreased with higher erosion of the NAWM mask) and the increase of both metrics with age, in line with others (**Figure 5**).^14, 49^

These observations have several implications in clinical and research settings. First, the close relationship between the FW and PET signals in this cohort challenges previous reports suggesting that the WM PET signal is myelin-dominated based on conventional FA. Our observations thus contribute to a better understanding and more accurate interpretation of the ambiguous WM signal in ^18^F-AV45 PET; this is of particular importance as Aβ-PET imaging is increasingly used in routine clinical practice for diagnostic workup of patients with suspect AD or mixed dementia, as well as for patient enrichment and target evaluation of anti-Aβ treatment trials. Second, in elderly subjects with suspicious areas of (low-to-moderate) cortical Aβ, who also have high WMH burden and atrophy, the cortical Aβ-PET signal may visually appear artificially reduced by the spill-in from low WM-associated signals, potentially leading to a false negative reading. On the other hand, in patients with substantial cortical Aβ and WMH, the cortex-WM contrast may visually appear enhanced (despite spill-over effects at the cortex-WM boundary). Third, although lower FW-adjusted FA did not translate into a lower SUVR signal in the *whole* NAWM, we found a lower SUVR in the NAWM of subjects with >30cc WMH volumes (**Supplementary Figure 7**). This may be due to microstructural changes spreading beyond the visible damage of the WMH (assessed on structural imaging) into the surrounding penumbras (peri-lesional areas) of WMH and connected tracts.^50, 53^ Ferris et al.^18^ recently showed that interstitial fluid accumulation, but not demyelination, extended up to 4mm beyond the boundaries of the WMH lesion – which could be in line with our lower NAWM PET signal in those with >30cc WMH (despite 2mm isotropic erosion of the NAWM mask). Thus, the spectrum of WM damage identified by PET may be wider than that identified by T1w or FLAIR sequences where more subtle microstructural changes may not manifest as contrast changes and can be hard to identify. Another important implication of the altered PET signal in WM, particularly to research applications, is that it will affect the use of WM as a reference region to quantify cortical Aβ SUVR. Future studies should mask out WMH from the WM reference region, erode their WM masks, and/or use WMH volume as a covariate in the analyses. They should also compare WM SUV between diagnostic groups to assure no significant differences in reference region uptake.^2, 54^

In relation to cognition, we observed that higher FW was associated with cognitive impairment both in regions of WMH and NAWM. The association in the NAWM may potentially reflect mild vascular damage and an early, pre-lesional role for FW in affecting cognitive function.^18, 26^ Apart from the significant FW-cognition relationship, we observed that FW (but not FW-adjusted FA) mediated the association between PET and executive function in WMH. In contrast, lower FW-adjusted FA only showed a weak relationship with lower MoCA score within WMH and with lower MMSE score within NAWM (the sensitivity of MoCA to WMH may be related to its wider application as a screening tool for vascular cognitive impairment by covering domains of executive function)^55^. Similarly, Maillard et al.^56^ found that FW fully mediated the effect of WMH volume on cognition among older adults, while they reported a lack of association between FW-adjusted FA and cognition. Taken together, these findings underscore FW as a potential biomarker in early dementia stages, following its appearance and clinical relevance in both pre-lesional and MRI-visible lesions of the WM. Thus, eliminating or preventing excessive FW in the WM may serve as a potential preventive strategy (e.g., through earlier, more personalized vascular risk control) against vascular injury, cognitive decline, and progression to AD dementia.

Limitations of our study include the use of single-shell and multi-center dMRI data. While our group performed great efforts to harmonize MRI acquisition parameters across different centers and cohorts, the dMRI data in MITNEC was acquired with fewer diffusion encoding directions than ADNI.^31^ Second, our study did not involve dynamic PET imaging with arterial input function. Thus, we cannot investigate whether WM PET may be affected by altered BBB permeability and/or the ligand’s pharmacokinetics through slower perfusion in WMH.^12, 13, 57^ Prior work with ^18^F-AV45 has shown that a 20% reduction in WM blood perfusion was associated with less than 5% reduction in SUVR at 50-60min post-injection;^58^ as such, we do not believe that that the ∼14% reduction in WMH SUVR (compared to NAWM) would be fully attributable to changes in blood perfusion. Another factor potentially affecting WM uptake may be related to the radioligand’s unique chemical and physical properties (e.g., lipophilicity).^59^ For those ligands that have been directly compared, WM retention has been highest for ^18^F-flutemetamol and was comparable between ^11^C-PiB and ^18^F-florbetapir.^60^ However, the topographic patterns of WM uptake (with lower uptake in WMH vs NAWM) was similar between ^18^F-flutemetamol and ^11^C-PiB irrespective of age.^61^ Future studies are needed to replicate our findings with different amyloid ligands in a mixed dementia cohort. Third, individuals were recruited from different cohorts/clinics. However, all analyses were controlled for demographic variables and were repeated within the cohorts separately, and further adjustment for this factor did not change the main results. Last, we focused on the averaged PET and DTI metrics within the NAWM and WMH. Thus, we cannot exclude that the FW-adjusted tissue compartment had more regional or tract-specific associations with Aβ-PET and cognitive performance.^26^

A major strength of our work is the inclusion of real-world patients covering the spectrum of low to severe WMH burden and cortical Aβ deposition. Specifically, seventy-eight subjects (∼70%) had WMH>10cc, allowing us to more accurately account for PVE which are prominent in most dementia studies with small WMH burden such as ADNI. In addition, by leveraging a novel dMRI technique that adjusts FA for FW contributions, as well as optimized structural pipelines for high WMH burden including a state-of-the-art deep learning segmentation model, we provided novel insights into the relationship of WM microstructure with the ^18^F-AV45 signal in the WM.

In conclusion, this study investigated the neurobiological underpinnings of Aβ-PET (^18^F-AV45) uptake in the WM – a marker of WM injury that is extracted “for free” in the diagnostic workup of dementia – using advanced diffusion modelling in a WMH enriched cohort. We found that lower ^18^F-AV45 PET uptake in MRI-visible WM lesions is strongly linked to elevated FW, potentially reflecting vascular damage, edema, inflammation, and/or proportional loss of myelin/WM tissue. On the other hand, in the NAWM, the ^18^F-AV45 PET uptake was more closely associated with Aβ of the cortex together with alterations to WM fiber integrity. We also highlighted that WM PET may be sensitive to microstructural changes in the penumbras surrounding WMH that are not yet visible on structural imaging. Last, in relation to cognition, higher FW both in WM lesions and NAWM related more closely to cognitive impairment than FW-adjusted FA or WM PET. Our study contributed to important new insights into the biological processes underlying the altered PET signal in the WM, further aiding in the interpretation of Aβ-PET studies. In addition, we present evidence that supports the need for accounting for WM lesions in Aβ-PET analyses. Finally, we highlighted FW as a promising and potentially early vascular-related injury marker in AD.

## Acknowledgements

We would like to express our deepest gratitude towards all the participants and caregivers for their support and participation in this study. We are grateful for the support of the Medical Imaging Trial Network of Canada (MITNEC; https://clinicaltrials.gov/ct2/show/NCT02330510?term=mitnec+sandra+black&rank=1) Grant #NCT02330510, and to Eli Lilly & Company for providing the 18F-florbetapir ligand. In addition, part of the data collection and sharing for this project was funded by ADNI (National Institutes of Health Grant U01 AG024904) and DOD ADNI (Department of Defense award number W81XWH-12-2-0012). ADNI is funded by the National Institute on Aging, the National Institute of Biomedical Imaging and Bioengineering, and through generous contributions from the following: AbbVie, Alzheimer’s Association; Alzheimer’s Drug Discovery Foundation; Araclon Biotech; BioClinica, Inc.; Biogen; Bristol-Myers Squibb Company; CereSpir, Inc.; Cogstate; Eisai Inc.; Elan Pharmaceuticals, Inc.; Eli Lilly and Company; EuroImmun; F. Hoffmann-La Roche Ltd and its affiliated company Genentech, Inc.; Fujirebio; GE Healthcare; IXICO Ltd.; Janssen Alzheimer Immunotherapy Research & Development, LLC.; Johnson & Johnson Pharmaceutical Research & Development LLC.; Lumosity; Lundbeck; Merck & Co., Inc.; Meso Scale Diagnostics, LLC.; NeuroRx Research; Neurotrack Technologies; Novartis Pharmaceuticals Corporation; Pfizer Inc.; Piramal Imaging; Servier; Takeda Pharmaceutical Company; and Transition Therapeutics. The Canadian Institutes of Health Research provided some funding to support ADNI clinical sites in Canada. Private sector contributions are facilitated by the Foundation for the National Institutes of Health (www.fnih.org). The grantee organization is the Northern California Institute for Research and Education, and the study is coordinated by the Alzheimer’s Therapeutic Research Institute at the University of Southern California. ADNI data are disseminated by the Laboratory for Neuro Imaging at the University of Southern California.

## Authors’ contributions

JO, MO, MG, and SEB designed the study and experiments. JO and MO wrote the manuscript, analyzed and interpreted the data with input from MG and SEB. MSK, KZ, PHK, WS, SA, CS, JR, AP, PM, and BL contributed to data generation, interpretation, and revision. JAC and OP contributed to toolbox development and revised the manuscript. AK, SS, CB, MB, HC, RF, RH, RJL, MDN, FSP, DJS, EES, VS, AT, JPS, and JCT participated in the study concept and design as site leaders and revised the manuscript. MG and SEB supervised all aspects of this work.

## Supplementary material

Supplementary material for this paper can be found at http://jcbfm.sagepub.com/content/by/supplemental-data

