## Supplementary materials for "Amyloid-PET of the white matter: relationship to free water, fiber integrity, and cognition in patients with dementia and small vessel disease"

** co-first authors,* † *co-senior authors*

**Table S1. Inclusion and Exclusion Criteria for MITNEC-C6.**

| **Inclusion Criteria** | **Exclusion Criteria** |
| --- | --- |
| 1. A. Early AD or amnestic, non‐amnestic single or multi-domain MCI with confluent pvWMH; recruited from memory clinics or B. minor stroke (e.g., subcortical lacunar infarct ≤1.5cm) or a TIA with confluent pvWMH; recruited from stroke prevention clinics 2. Age ≥ 60 3. Written informed consent 4. More than 8 years of education 5. Expected survival greater than 2 years 6. Sufficiently fluent in French or English for cognitive testing 7. Mini-Mental State Exam score ≥ 20 8. pvWMH score on CT or MRI of ≥2 on the periventricular Fazekas scale* | 1. Cortical or non-lacunar infarct on imaging 2. Persisting hemiparesis after a motor stroke, leg strength <4/5 on the Medical Research Council (MRC) scale; significant cerebellar ataxia 3. Contraindications to 3T MRI 4. Major psychiatric disorder in the preceding 5 years 5. History of substance abuse in the past 2 years 6. Serious/chronic systemic or neurological illness (other than AD) such as Parkinson’s disease, multi-infarct dementia, Huntington’s disease, normal pressure hydrocephalus, brain tumor, progressive supranuclear palsy, seizure disorder, subdural hematoma, multiple sclerosis, or history of significant head trauma followed by persistent neurologic deficits or structural brain abnormalities 7. Pain or sleep disorder that could interfere with testing 8. Claustrophobia 9. Radiation therapy to the head or neck or in a research study involving radiation 10. Unable or unwilling to comply with protocol requirements or deemed by the investigator to be unfit for the study |

* Fazekas 2 can be included if bilateral posterior or anterior periventricular caps extending at least 10mm from the ventricle (i.e., halfway into surrounding white matter vs. extending out to most surrounding white matter for Fazekas 3)

**Table S2. Description of MR imaging parameters.**

| *Study* | MITNEC – C6 | | |
| --- | --- | --- | --- |
| Sequence | 3DT1 | | |
| ***Protocol*** | | | |
| Vendor | GE | Philips | Siemens |
| Field Strength | 3T | 3T | 3T |
| Model | Discovery | Achieva | Skyra/Trio/Prisma |
| Version | 22 | 3.2.3 |  |
| Sequence Name | 3D FAST SPGR | 3D TFE | 3D MP-RAGE |
| Imaging Options | IrP- Asset | Fast (Sense) | iPat |
| ***Pulse Timing*** | | | |
| TE (ms) | Min full | Min (3.3) | 2.98 |
| TR (ms) | Min | Min (7.3) | 2300 |
| Flip Angle (°) | 11 | 9 | 9 |
| TI (ms) | 400 | 945 | 900 |
| **Scan Range** | | | |
| FOV (in-plane) (mm) | 256 x 256 | 256 x 248 | 256 x 256 |
| Slice Thickness (mm) | 1 | 1 | 1 |
| Gap Between Slices (mm) | 0 | 0 | 0 |
| No. Slices | 176 | 176 | 176 |
| **Acquisition** | | | |
| Orientation | Sagittal | Sagittal | Sagittal |
| Matrix Size | 256 x 256 | 256 x 248 | 256 x 256 |
| Voxel Size [L/R x A/P x I/S] | 1 x 1 x 1 | 1 x 1 x 1 | 1 x 1 x 1 |
| NEX | 1 | 1 | 1 |
| Acceleration Factor (Parallel factor) | 2 | 2 | 2 |
| **Other** | | | |
| Fat Suppression | None | None | None |
| Bandwidth | 31.25 (kHz) | 228 (Hz/px) | 240 (Hz/px) |
| Echo Train Length | - | - | - |
| **Coil Type** | | | |
| Head | X | X | X |
| Channel | 8-12 (HNS) | 8 | 12(Trio) 20 (Prisma) |

| Sequence | 2D FLAIR | | |
| --- | --- | --- | --- |
| ***Protocol*** | | | |
| Sequence Name | 2D T2FLAIR | 2D IR TSE | 2D IR TDF |
| Imaging Options | EDR, IR | Fast (Sense) | iPat |
| ***Pulse Timing*** | | | |
| TE (ms) | 140 | 125 | 120 |
| TR (ms) | 9000 | 9000 | 9000 |
| Flip Angle (°) | 125 | 90 (150 refocus) | 165 |
| TI (ms) | 2250 | 2500 | 2500 |
| **Scan Range** | | | |
| FOV (in-plane) (mm) | 240 x 240 | 240 x 240 | 240 x 240 |
| Slice Thickness (mm) | 3 | 3 | 3 |
| Gap Between Slices (mm) | 0 | 0 | 0 |
| No. Slices | 48 | 48 | 48 |
| **Acquisition** | | | |
| Orientation | Oblique Axial | Oblique Axial | Oblique Axial |
| Matrix Size | 256 x 256 | 256 x 242 | 256 x 256 |
| Voxel Size [L/R x A/P x I/S] | 0.94 x 0.94 x 3 | 0.94 x 0.99 x 3 | 0.94 x 0.94 x 3 |
| NEX | 1 | 1 | 1 |
| Acceleration Factor (Parallel factor*) | No Asset | 2 (SENSE) | 2 |
| **Other** | | | |
| Fat Suppression | None | None | None |
| Bandwidth | 25 (kHz) | 242 (Hz/px) | 220 (Hz/px) |
| Echo Train Length |  | 19 | 19 |
| **Coil Type** | | | |
| Head | X | X | X |
| Channel | 8-12 (HNS) | 8 | 12(Trio) 20 (Prisma) |

| *Sequence* | DIFFUSION MRI | | |
| --- | --- | --- | --- |
| ***Protocol*** | | | |
| Vendor | GE | Philips | Siemens |
| Field Strength | 3T | 3T | 3T |
| Model | Discovery | Achieva | Skyra/Trio/Prisma |
| Version | 22 | 3.2.3 |  |
| Sequence Name | DWI | DWI | DWI |
| Imaging Options | ASSET | Sense | iPat |
| ***Pulse Timing*** | | | |
| TE (ms) | Min | 100 | 96/ 63(Prisma) |
| TR (ms) | 9000 | 9931 | 9400 |
| Flip Angle (°) | 90 | 90 | 90 |
| TI (ms) | - | - | - |
| ***Scan Range*** | | | |
| FOV (in-plane) (mm) | 256 x 256 | 256x 256 | 256 x 256 |
| Slice Thickness (mm) | 2 | 2 | 2 |
| Gap Between slices (mm) | 0 | 0 | 0 |
| No. Slices | 70 | 70 | 70 |
| ***Acquisition*** | | | |
| Orientation | Oblique Axial | Oblique Axial | Oblique Axial |
| Matrix Size | 128 x 128 | 128 x 128 | 128 x 128 |
| Voxel Size (L/R x A/P x I/S) | 2 x 2 x 2 | 2 x 2 x 2 | 2 x 2 x 2 |
| NEX | 1 | 1 | 1 |
| Acceleration Factor (Parallel factor*) | 2 | 2 | 2 |
| b-value 1 | 0 | 0 | 0 |
| b-value 2 | 1000 | 1000 | 1000 |
| Number of Directions | 30 | 32 | 30 |
| ***Other*** | | | |
| Fat Suppression | FatSat | FatSat | FatSat |
| Bandwidth (Hz/Px) |  | 2045 | 2056 |
| T2 Images | 3 | 3 | 3 |
| EPI Factor |  | 67 | 128 |
| Gradients |  |  | Monopolar (Prisma) |
| ***Coil Type*** | | | |
| Head | x | x | x |
| Channel | 8-12 (HNS) | 8 | 12(Trio) 20 (Prisma) |

ADNI-2: <http://adni.loni.usc.edu/wp-content/uploads/2010/05/ADNI2_GE_3T_22.0_T2.pdf>

**Table S3.** Relationship between ^18^F-AV45 SUVR and DTI metrics in regions of the WMH vs NAWM with additional adjustment for cohort (ADNI vs MITNEC) across all subjects and for clinic (stroke prevention vs dementia clinic) in the high WMH group.

| All subjects (n=115) | | | | | | |
| --- | --- | --- | --- | --- | --- | --- |
| **SUVR** | **FW** | | | **FW-adjusted FA** | | |
|  | β | *P* | 95%CI_bs_ | β | *P* | 95%CI_bs_ |
| WMH | -0.52 | ***0.009*** | -0.86,-0.18 | -0.05 | *0.61* | -0.24,+0.13 |
| NAWM | -0.21 | *0.058* | -0.41,+0.06 | -0.16 | *0.064* | -0.34,+0.03 |
| High WMH group (n=58) | | | | | | |
| **SUVR** | **FW** | | | **FW-adjusted FA** | | |
|  | β | *P* | 95%CI_bs_ | β | *P* | 95%CI_bs_ |
| WMH | -0.33 | ***0.022*** | -0.56,-0.06 | -0.16 | *0.27* | -0.40,+0.14 |
| NAWM | -0.16 | *0.24* | -0.41,+0.12 | -0.30 | ***0.011*** | -0.52,-0.05 |

**Table S4.** Relationship between DTI metrics and cognition in regions of the WMH vs NAWM with additional adjustment for clinic (stroke prevention vs dementia) in the high WMH group.

| **Regions of white matter hyperintensities** | | | | | | |
| --- | --- | --- | --- | --- | --- | --- |
| High WMH group (n=58) | | | | | | |
|  | **FW** | | | **FW-adjusted FA** | | |
|  | β | *P* | 95%CI_bs_ | β | *P* | 95%CI_bs_ |
| Semantic | -0.25 | ***0.067*** | -0.56,+0.03 | +0.08 | *0.53* | -0.13,+0.34 |
| Language | -0.40 | ***0.003*** | -0.69,-0.15 | +0.23 | ***0.083*** | -0.05,+0.46 |
| MoCA | -0.33 | ***0.013*** | -0.58,+0.03 | +0.12 | *0.35* | -0.12,+0.37 |
| MMSE | -0.30 | ***0.041*** | -0.55,-0.03 | +0.08 | *0.58* | -0.16,+0.40 |
| Speed | +0.07 | *0.61* | -0.22,+0.36 | -0.11 | *0.41* | -0.33,+0.12 |
| Executive | +0.20 | *0.16* | -0.04,+0.51 | -0.18 | *0.19* | -0.52,+0.07 |
| **Region of normal-appearing white matter** | | | | | | |
| High WMH group (n=58) | | | | | | |
|  | **FW** | | | **FW-adjusted FA** | | |
|  | β | *P* | 95%CI_bs_ | β | *P* | 95%CI_bs_ |
| Semantic | -0.37 | ***0.013*** | -0.67,-0.07 | -0.09 | *0.55* | -0.33,+0.21 |
| Language | -0.24 | *0.11* | -0.55,+0.03 | +0.15 | *0.33* | -0.10,+0.53 |
| MoCA | -0.26 | ***0.067*** | -0.61,+0.02 | +0.07 | *0.65* | -0.18,+0.37 |
| MMSE | -0.34 | ***0.030*** | -0.68,-0.06 | +0.13 | *0.42* | -0.17,+0.47 |
| Speed | +0.21 | *0.17* | -0.10,+0.50 | +0.10 | *0.49* | -0.16,+0.40 |
| Executive | +0.14 | *0.35* | -0.14,+0.42 | -0.10 | 0.50 | -0.40,+0.10 |

Following cognitive tests were applied. Semantic fluency: animal naming; Language: BNT; Speed: TMT-A; Executive function: TMT-B

**Methods S1. Structural MRI.** Description of WMH volume extraction.

To segment WMH volumes, we employed our in-house developed HyperMapper tool (<https://hypermapp3r.readthedocs.io/>), which outperforms state-of-the-art segmentation techniques.^1^ HyperMapper is based on a novel Bayesian 3D convolutional neural network (CNN) with a U-Net architecture that automatically segments WMH and provides uncertainty estimates of the segmentation output for quality control. The CNN was trained using 432 subjects recruited from four multisite imaging studies with different augmentation schemes including the addition of noise, various resolutions, and changes in contrast, to make the model more robust to the challenges that commonly result from different MRI scanners and acquisition protocols. It can deal with brains with extensive atrophy and segments the WMH in seconds. More details can be found in Mojiri et al.^1^ HyperMapper uses T1-weighted MRI, FLAIR (co-registered to T1w using affine registration), and a brain mask as the inputs.

**Methods S2. Diffusion Tensor Imaging (DTI).** Description of DTI processing methods.

The diffusion-weighted MRI (dMRI) data were processed using FSL (<http://www.fmrib.ox.ac.uk/fsl>), MRtrix3 (<https://www.mrtrix.org/>), and ANTS (http://stnava.github.io/ANTs/). For each subject, the single *b*0 images were skull stripped using FSL *bet* and co-registered to T1w using affine registration with ANTS. The dMRI images were bias-corrected using MRtrix *dwibiascorrect* and transformed to T1w space. They were denoised using the MRtrix *dwidenoise*. Head motion and eddy-current induced distortion were corrected using MRtrix *eddy_correct*. Images were visually inspected for excessive motion, signal drop-out, and/or artefacts (*N*=5 were removed, leaving N=115 total). Tensor (ellipsoid) fitting was conducted with a weighted-least square at each brain voxel using FSL *dtifit*. The resulting eigenvalues (λ_1_, λ_2_, λ_3_), capturing the dimensions of each voxel’s ellipsoid fit, were used to create the DTI scalar metrics. These metrics include fractional anisotropy (FA) and mean diffusivity (MD). High FA is thought to arise from higher diffusivity along one preferred direction (i.e., along intact myelinated fibers, thus representing fiber integrity) or from a loss of crossing fibers.^2,3^ On the other hand, MD represents the magnitude of diffusion (average of eigenvalues) regardless of the directionality.

For FW mapping, the eddy current and motion-corrected dMRI data were fitted to a two-compartment diffusion model in each voxel, separating the FW compartment from the non-FW tissue compartment.^4^ Specifically, the isotropic compartment has a fixed diffusion coefficient (3×10^−3^mm^2^/s) and models the freely diffusing extracellular water molecules (free water, FW). On the other hand, the tissue-specific compartment models the water molecules that are close to the cellular membranes of brain tissue (restricted diffusion) and is presented by the diffusion tensor. The FW was scaled by the signal acquired for zero diffusion weightings,^4^ while FA, MD, and FW-adjusted FA were scaled by their respective whole brain values assuming site/scanner differences may affect the global metric.^5,6^

**Methods S3. Amyloid-PET imaging.** Description of PET processing methods.

The PET protocol stipulated that 370MBq of ^18^F-AV45 was administered, followed by 20min dynamic scanning at 50min post-injection, and scatter and attenuation correction. The individual frames (4x5min) were motion-corrected to the first frame and averaged to generate one static image. This image was then co-registered to T1w MRI using PetSurfer (version 6.0) *mri_coreg*. The images were smoothed to a common Gaussian kernel of 8mm FWHM across sites, and the grey matter values were corrected for partial volume effects using GTM implemented in PetSurfer *mri_gtmpvc*. A quantitative cut-off SUVR=1.1 (not partial volume corrected) was determined from the AD-signature regions using Gaussian mixture modeling of our cohort.^7^

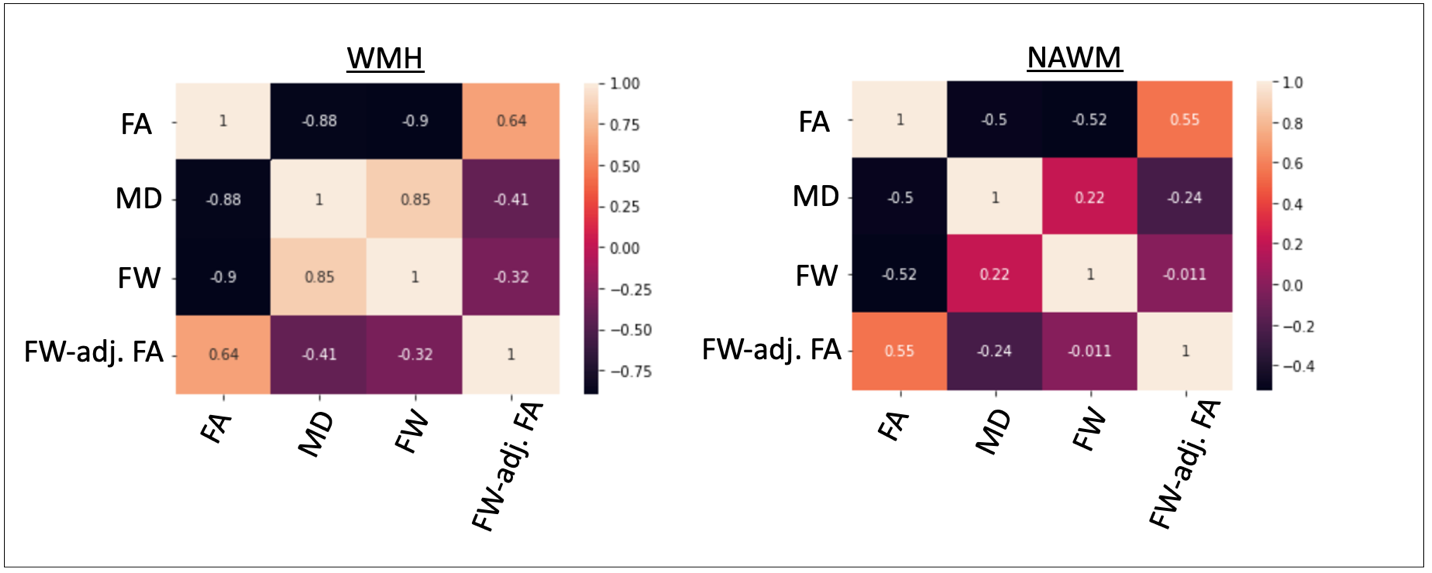

**Figure S1**. Correlation coefficients (Pearson’s R) between DTI metrics across all subjects. Abbreviations: FA, fractional anisotropy; FW, free water; MD, mean diffusivity; NAWM, normal appearing white matter; WMH, white matter hyperintensities

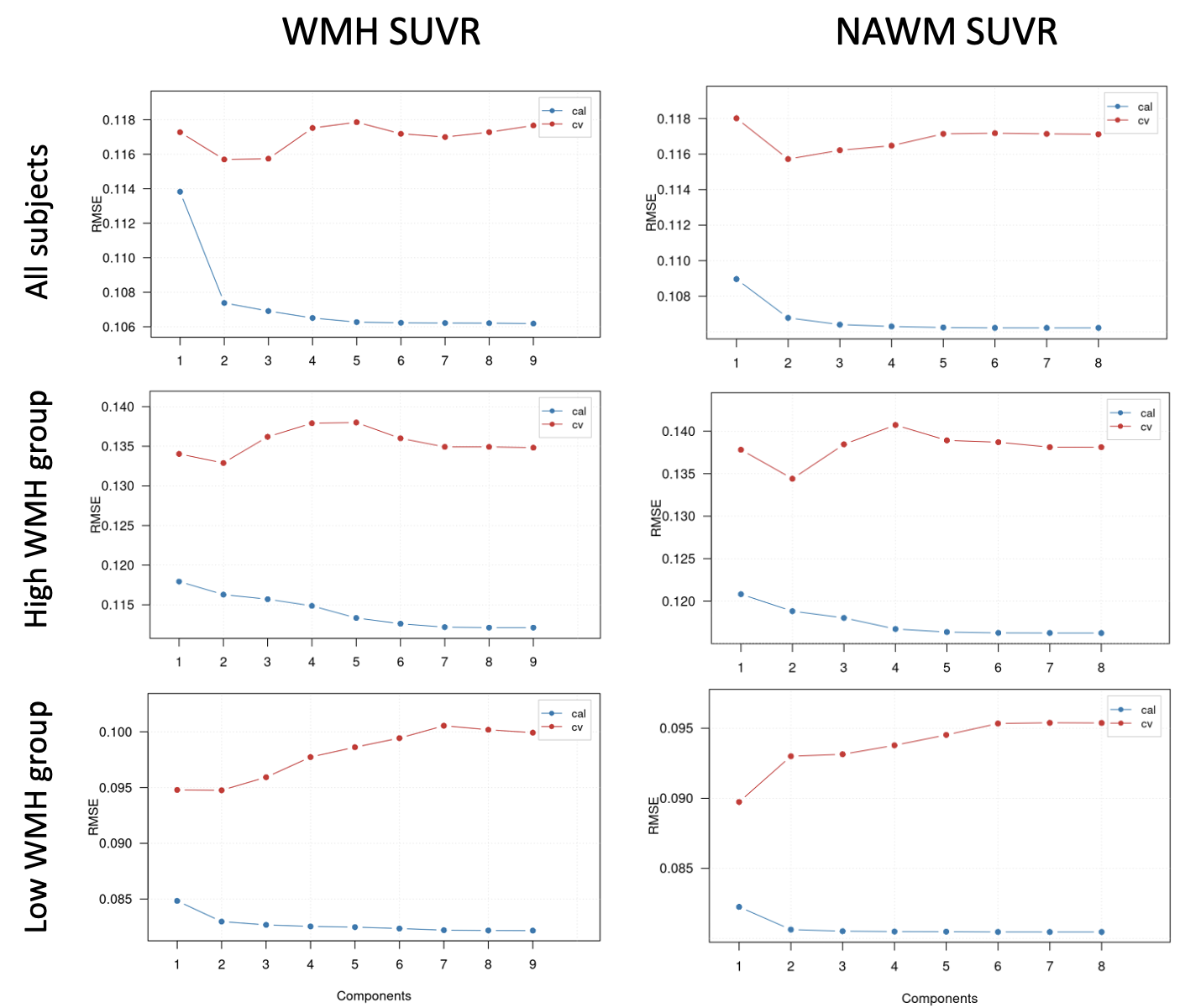

**Figure S2.** Results of the cross-validation of the partial-least-squares regression model. Root mean squared error (RMSE) of predictor curves for the calibrated (cal; blue) and the cross-validated (cv; red) predictions of the WM SUVR. The minimum RMSE(cv) corresponds to the optimal amount of selected components (one or two, depending on the cohort).

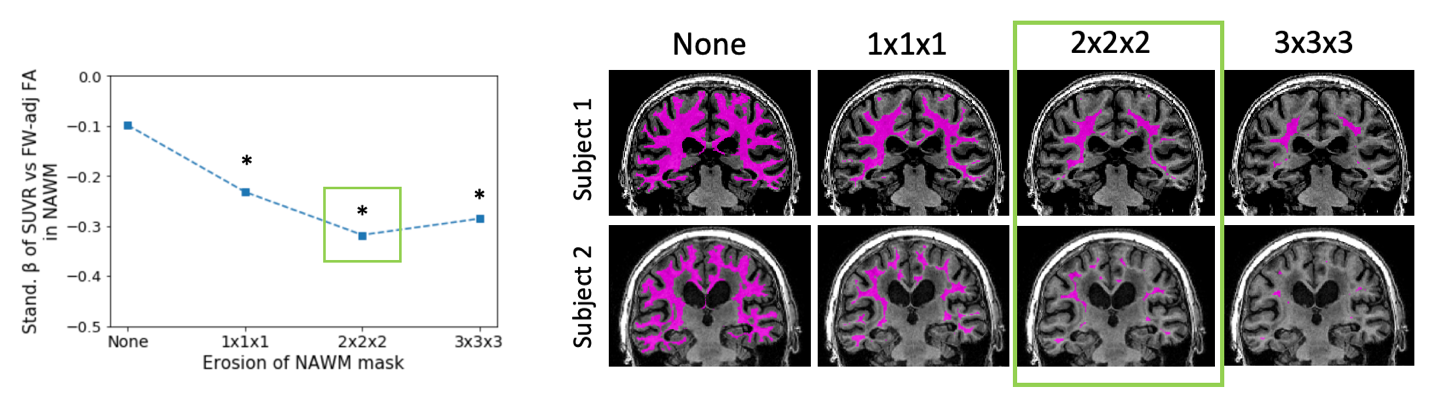

**Figure S3**. Influence of different NAWM erosion kernels. (Left) Standardized β of the relationship between FW-adjusted FA and SUVR in NAWM for the different erosion kernels (none, 1x1x1, 2x2x2, and 3x3x3 mm^3^) in the high WMH group; stars indicate significance (P<0.05 with adjustment for the covariates). (Right) Examples of the NAWM mask after erosion for two representative subjects (subject 1 and 2: total WMH volume = 16.00 and 67.29 cc, respectively). Abbreviations: FW-adj FA, free water-adjusted fractional anisotropy; NAWM, normal appearing white matter; SUVR, standardized uptake value ratio

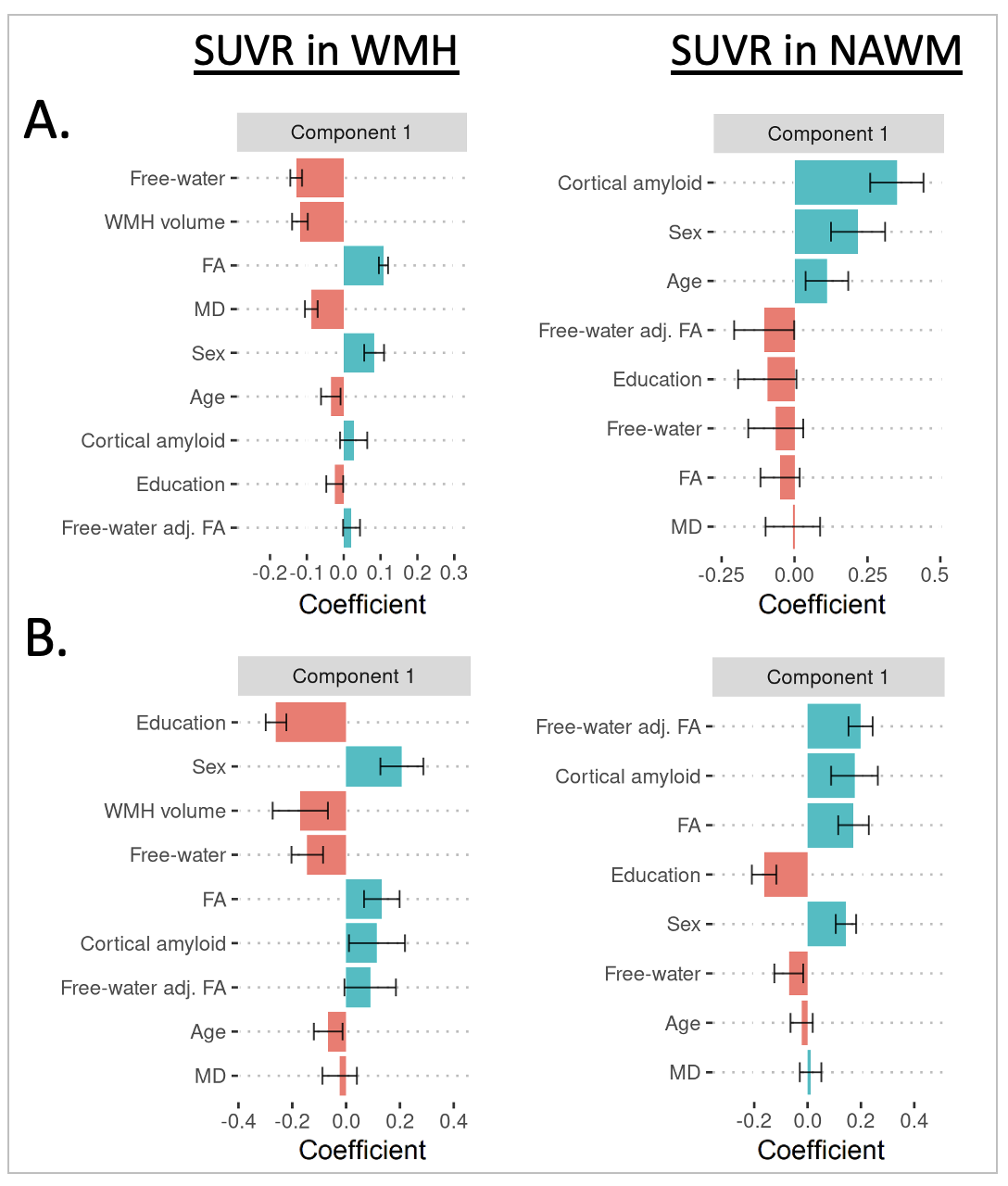

**Figure S4**. PLS analysis across all subjects (panel A) and within the low WMH group (panel B) showing the relationship of ^18^F-AV45 SUVR with DTI and demographical/imaging variables in a) WMH (component 1 ~ 18%, component 2 [data not shown] ~ 9%) and b) NAWM (component 1 ~ 24%, component 2 [data not shown] ~ 3%). Error bars represent 95%CI based on bootstrapping with 5,000 repetitions. Abbreviations: FW, free water; MD, mean diffusivity; NAWM, normal appearing white matter; SUVR, standardized uptake value ratio; WMH, white matter hyperintensities

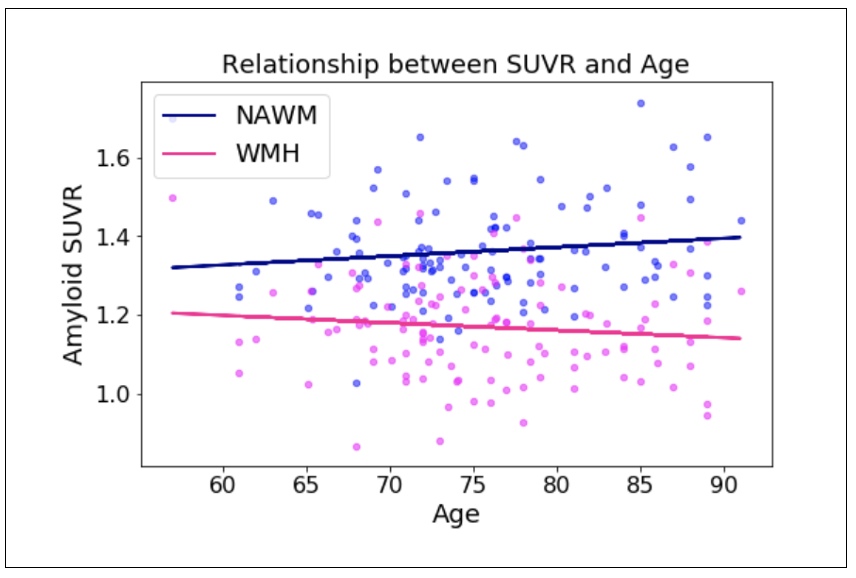

**Figure S5.** Interaction effect (P<0.0001) between age and WM region (NAWM vs. WMH) on amyloid SUVR. Abbreviations: NAWM, normal appearing white matter; SUVR, standardized uptake value ratio; WMH, white matter hyperintensities

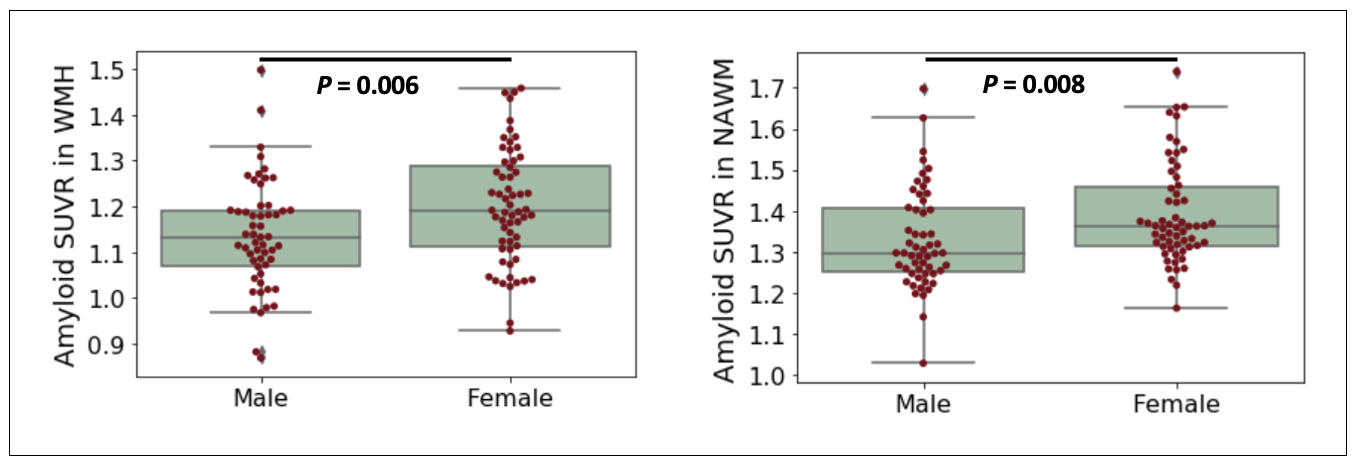

**Figure S6**. Females showed higher amyloid SUVR in WMH (left) and NAWM (right) compared to males. Abbreviations: NAWM, normal appearing white matter; SUVR, standardized uptake value ratio; WMH, white matter hyperintensities

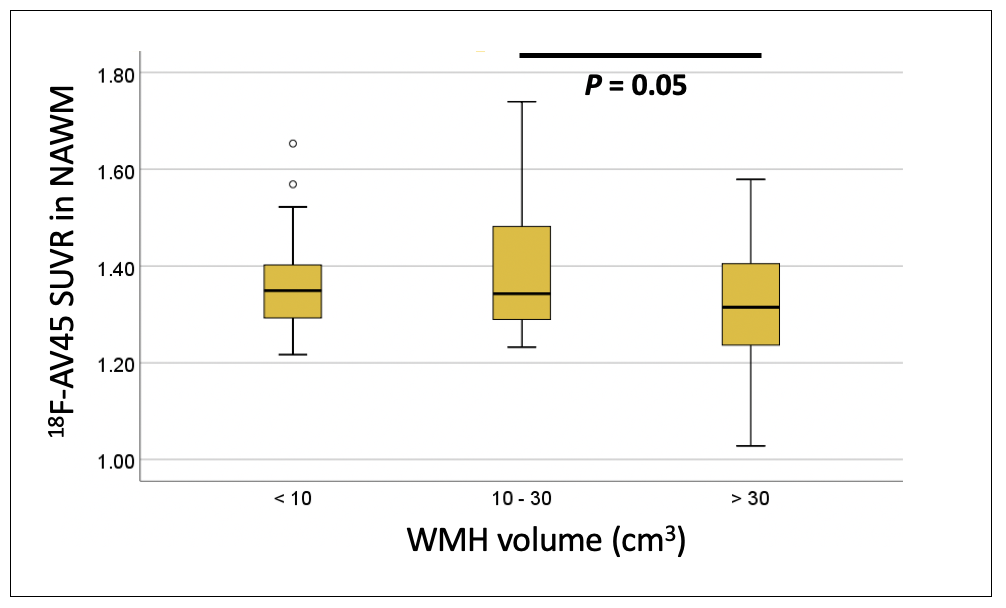

**Figure S7**. ^18^F-AV45 SUVR in NAWM is significantly lower in subjects with high WMH volume (>30cc). Significance is based on ANOVA with Tukey multiple comparisons test.
